# Lin28a rejuvenates muscle stem cells via mitochondrial optimization

**DOI:** 10.1101/2021.10.14.462144

**Authors:** Peng Wang, Xupeng Liu, Jun-Hao Elwin Tan, Min-Wen Jason Chua, Yan-Jiang Benjamin Chua, Lanfang Luo, Shilin Ma, Wenhua Cao, Wenwu Ma, Ziyue Yao, Yu Chen, Hefan Miao, Luyao Guo, Liping Zhang, Lu Guang, Kun Liang, Yuefan Wang, Jiali Su, Shuqing Liu, Ruirui Liu, Ruiqi Rachel Wang, Chunwei Li, Na Ai, Yun Li, Zongming Jiang, Taoyan Liu, Bin Tean Teh, Lan Jiang, Kang Yu, Ng Shyh-Chang

## Abstract

The well-conserved correlation between juvenility and tissue regeneration was first discussed by Charles Darwin. Ectopic Lin28 is known to play an important role in somatic reprogramming and tissue regeneration, but endogenous Lin28’s role in tissue homeostasis and juvenility had remained unclear. Through lineage tracing, we found that a rare subset of muscle stem cells (MuSCs) expressing Lin28a can respond to acute injury by proliferating as Pax3+ or Pax7+ MuSCs, and contribute to all types of myofibers during muscle regeneration. Compared with conventional Pax7+ MuSCs, Lin28a+ MuSCs express more Pax3 and show enhanced myogenicity in vitro. In terms of the epigenetic clock, adult Lin28a+ MuSCs lie between adult Pax7+ MuSCs and embryonic Pax7+ myoblasts according to their DNA methylation profiles. We found that Lin28a+ MuSCs upregulate several embryonic limb bud mesoderm transcription factors and could maintain a juvenile state with enhanced stem cell self-renewal and stress-responsiveness in vitro and in vivo. When combined with telomerase and *TP53* inhibition to biomimic mouse Lin28a+ MuSCs, we found that Lin28a can rejuvenate and dedifferentiate aged human primary myoblasts into engraftable, self-renewing MuSCs. Mechanistic studies revealed that Lin28a activated the HIF1A pathway by optimizing mitochondrial ROS (mtROS), thereby rejuvenating MuSC self-renewal and muscle regeneration. Our findings connect the stem cell factor Lin28, mtROS metabolism and stress response pathways to the process of stem cell rejuvenation and tissue regeneration.

## Introduction

Despite enormous strides in our understanding of pluripotency, somatic cell reprogramming and transdifferentiation, the molecular basis for self-renewal of stemness has remained unclear. While we have succeeded in fully resetting somatic cells back to the pluripotent state, we have not been able to rejuvenate somatic cells by dialing back biological and/or epigenetic aging. Lin28 was one of the first genes discovered to be linked to developmental timing and juvenile programs of development, a concept related to stem cell self-renewal^1–3^, and which can be partly explained by the Lin28-*let-7* pathway^4–8^. In terms of the epigenetic clock for biological aging, a rejuvenation event was recently observed during mammalian pre-gastrulation embryogenesis^9^, when Lin28a is most highly expressed. In nematodes, lin-28 is also highly expressed during early development, especially in the hypodermis and muscle cells, but gradually diminishes and disappears by adulthood. Loss-of-function in lin-28 accelerates cellular differentiation, while gain-of-function in lin-28 promotes self-renewal for a few cell divisions^10^. Furthermore, mammalian studies have revealed that ectopic Lin28a overexpression promotes tissue regeneration^11^, but its endogenous role in mammalian tissue homeostasis had remained unclear.

Skeletal muscles possess strong regenerative abilities. Skeletal muscle satellite cells (MuSCs) are a group of resident stem cells, embedded between the muscle fibrillar membrane and the basement membrane, which undergo self-renewal and are necessary for skeletal muscle regeneration^12^. In injured muscles, MuSCs are activated and begin to proliferate as committed myogenic progenitor cells^13^. These activated myogenic cells fuse with existing myofibers, or form de novo myofibers, to achieve muscle repair and regeneration^14^. Although the upregulation of endogenous Lin28a has only been observed during muscle regeneration^15^, amongst all the normal adult tissues of mammals, it was unclear what cell-type(s) expressed Lin28a and whether Lin28a played a functional role in the maintenance of MuSC and/or myoblast self-renewal. This is partially because no rigorous lineage-tracing studies have been performed with Lin28a.

Here we took into account the fact that Lin28a mRNA is regulated post-transcriptionally^16^, and tagged Lin28a protein with a “self-cleaving” CreERT2 recombinase to trace Lin28a+ cell lineages in adult mouse muscles. To our surprise, we found a previously unknown and rare group of MuSCs that could form all types of muscle fibers in adult muscle, with characteristics of fetal limb muscle progenitors. Lin28a+ MuSCs manifest higher dedifferentiation, stress-responsiveness, self-renewal and myogenic capacity. Biomimicry of mouse Lin28a+ MuSCs, with their expression of Lin28a, the telomerase complex component Tep1 and the p53 inhibitor Mdm4, allowed us to rejuvenate aged human myoblasts into engraftable, self-renewing MuSCs. Mice inserted with an injury-responsive Lin28a transgene also showed that Lin28a promotes skeletal muscle regeneration by enhancing adult myoblast self-renewal in vivo. Mechanistically, we found that Lin28a operated independently of *let-7* microRNAs and enhanced oxidative metabolism by regulating the mRNAs of mitochondrial OxPhos enzymes, thereby optimizing mtROS, and activating a HIF1A-driven hypoxic and glycolytic stress response to rejuvenate adult myoblasts.

## Results

### Lin28a+ satellite cells contribute to all types of myofibers during regeneration

In order to observe whether Lin28a is expressed in a particular subset of muscle cells, we tagged Lin28a protein at its C-terminus with a “self-cleaving” T2A-CreERT2 using homologous recombination, constructed Lin28a-T2A-CreERT2 transgenic mice (Figures S1A,B), and bred the transgenic mice with Rosa26-loxp-stop-loxp-tdTomato (tdTO) reporter mice to trace the fate of Lin28a+ cells after muscle injury. After about 20 days of tracing (Figure 1A), we found that uninjured mice’ tibialis anterior (TA) muscles had a small amount of Lin28a-tdTO+ muscle fibers and Lin28a-tdTO+ mono-nuclear cells in the muscle interstitium (Figure 1B), suggesting that Lin28a+ cells existed in the muscle interstitium, and could fuse with ~10% of the existing TA muscle fibers during normal muscle homeostasis (Figure 1B). Upon injury, the numbers of Lin28a-tdTO+ muscle fibers and Lin28a-tdTO+ mono-nuclear cells were significantly increased (Figures 1B,C), suggesting that the injury stimulated Lin28a+ cell proliferation and increased fusion with existing myofibers by over 3-fold (Figure 1C). In another muscle group with slower turnover (soleus muscles)^17^, the number of Lin28a-tdTO+ myofibers was also significantly increased after injury (Figures 1B,C). To ensure our Lin28a-tdTO+ lineage tracing was specific and faithful to endogenous Lin28a expression, and given the sub-optimal performance of Lin28a antibodies in immunofluorescence sections, we examined the distribution of tdTO+ cells in the testis, where only spermatogonial stem cells were expected to express Lin28a ^18^. Indeed, we found that only a subset of PLZF+ spermatogonial stem cells on the periphery of seminiferous tubules were specifically labeled as Lin28a-tdTO+ cells after 20 days of lineage-tracing (Figure S1C). When Lin28a+ cells were cultured in vitro, the expression of both Lin28a protein and Cre protein were extinguished simultaneously, indicating that the turnover of “cleaved” T2A-CreERT2 was sufficiently rapid for it to accurately mark Lin28a+ cells with high fidelity (Figure S1D). To ensure the expression of endogenous Lin28a was not significantly interfered by the T2A-CreERT2, we examined total Lin28a protein expression in the muscles and testes of heterozygous Lin28a-T2A-CreERT2 mice, and found no significant differences between wild type and Lin28a-T2A-CreERT2; LSL-tdTO mice (Figure S1E).

**Figure 1.**
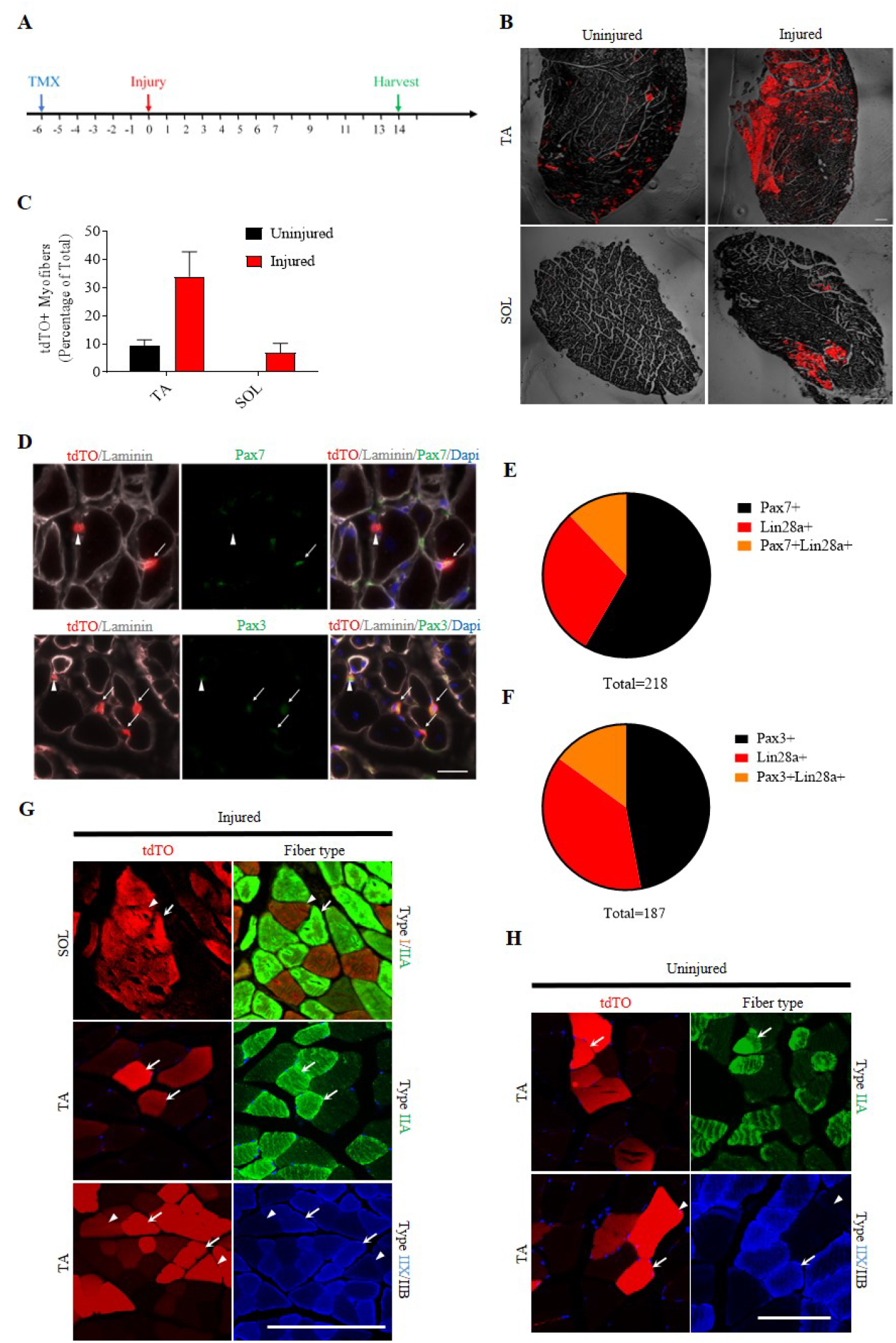
Lin28a+ satellite cells contribute to all types of myofibers during regeneration. **A**) Schematic of tamoxifen (TMX) treatment. Lin28a-T2A-CreER;LSL-tdTO mice were injected with TMX, 6 days before cryoinjury and daily for the first one week, every other day from the 7^th^ day after injury, and harvested on the 14^th^ day after injury. (**B**) Tibialis anterior (TA) and soleus (SOL) muscles of Lin28a-T2A-CreER;LSL-tdTO mice were injured by cryoinjury and harvested on day 14 post injury. CTL: contralateral tibialis anterior (TA) or soleus (SOL) muscle. Scale bar: 200μm. (**C**) Quantification of the number of tdTomato^+^ (tdTO+) muscle fibers per 100 muscle fibers in the TA and soleus muscles of Lin28a-T2A-CreER;LSL-tdTO mice on day 14 post injury and without injury. Quantification was performed on the images in Fig. 1C. For each muscle section, at least 3 whole sections were quantified and averaged. (**D**) TA muscle of Lin28a-T2A-CreER;LSL-tdTO mice injured by cryoinjury and harvested on day 14 post injury. Muscle sections were co-stained for laminin (gray), DAPI (blue) and Pax7 (green) or Pax3 (green). Arrows indicate Lin28a+Pax7+ or Lin28a+Pax3+ cells, and arrowheads indicate Lin28a+Pax3- or Lin28a+Pax7-cells. Scale bar: 20μm. (**E**) Quantification of the number of Pax7+, Lin28a+, and Pax7+/Lin28a+ double positive cells per field in TA muscle of Lin28a-T2A-CreER;LSL-tdTO mice on day 14 post-injury. Quantification was performed on the images in Fig. 1E. For each muscle section, at least 5 different fields were quantified and averaged. (**F**) Quantification of the number Pax3+, Lin28a+, and Pax3+/Lin28a+ double positive cells per field in TA muscle of Lin28a-T2A-CreER;LSL-tdTO mice on day 14 post-injury. Quantification was performed on the images in Fig. 1E. For each muscle section, at least 5 different fields were quantified and averaged. (**G**) Immunostaining of transverse sections of injured SOL and TA muscles obtained from Lin28a-T2A-CreER;LSL-tdTO mice at 14 days post-injury, for type I (red), type IIA (green), type IIX (blue) and type IIB (black) myofibers, compared to tdTO+ fluorescence (arrowheads). Scale bar: 100μm. (**H**) Immunostaining of transverse sections of uninjured TA muscles obtained from Lin28a-T2A-CreER;LSL-tdTO mice at day 14 of lineage tracing, for type IIA (green), type IIX (blue) and type IIB (black) myofibers, compared to tdTO+ fluorescence (arrowheads). Scale bar: 100μm.

Pax7 is generally regarded as the definitive marker of adult muscle stem cells (MuSCs)^19^, but reports have also shown that some MuSCs do not express Pax7, but express Pax3 instead^20–22^. Therefore, we sought to determine whether Lin28a-tdTO+ mono-nuclear cells expressed Pax3 or Pax7. Immunofluorescence results showed that all the Lin28a-tdTO+ mono-nuclear cells were located between the basement membrane and myofibrillar membrane, similar to the position of Pax7+ muscle satellite cells, but only a fraction of them co-expressed Pax7 or Pax3 (Figure 1D). Overall, Lin28a-tdTO+ cells only made up <30% of the Pax7+ or Pax3+ MuSC pool (Figures 1E,F), suggesting they constituted a minor fraction of the Pax7/Pax3+ MuSC population. Quantification showed that 37.1% of Lin28a-tdTO+ mono-nuclear cells co-expressed Pax7 (Figure 1E), whereas 40.6 % of Lin28a-tdTO+ mono-nuclear cells co-expressed Pax3 (Figure 1F), and the remainder of Lin28a-tdTO+ cells were Pax7-Pax3-. Overall, our Western blot and lineage tracing results suggested that at least some of the satellite cells or MuSCs express Lin28a but not Pax7 nor Pax3, and these Lin28a+ MuSCs can respond to injury by transiently proliferating as Lin28a-Pax7+/Pax3+ MuSCs, thereby contributing to skeletal muscle regeneration.

Skeletal myofibers show a certain level of diversity in metabolic and physical function upon terminal differentiation, with each muscle group containing a mixture of different types of muscle fiber fates, e.g. type I, IIa, IIx, and IIb myofibers. These myofibers can be roughly classified as slow-twitch (I) and fast-twitch (IIa, IIx, IIb), or oxidative (I, IIa) and glycolytic (IIx, IIb)^23^. Given the difference in Lin28a-tdTO+ labeling efficiency in fast-twitch TA and slow-twitch soleus muscles, we sought to determine whether the types of muscle fibers formed by Lin28a-tdTO+ cells are specific. Immunofluorescence staining of uninjured mouse muscle fibers showed that during the 20-day tracing window, Lin28a-tdTO+ cells can contribute to type IIa, IIx, and IIb muscle fibers in TA muscles (Figure 1H), but not the type I myofibers in soleus muscles (Figures 1B,C). However, after injury, Lin28a-tdTO+ cells could contribute to all types of muscle fibers in both the slow-twitch soleus and fast-twitch TA muscles (Figure 1G). These results suggest that Lin28a+ MuSCs can contribute to all types of muscle fibers after proliferation and differentiation during skeletal muscle regeneration in vivo.

### Lin28a+ cells are MuSCs which show enhanced myogenic potential in vitro

In order to further determine the properties of Lin28a-tdTO+ mono-nuclear cells, relative to conventional Pax7+ MuSCs, we decided to use flow cytometry to analyze this group of cells. On the tdTomato channel, we observed that tdTO+ cells increased significantly only after muscle injury (P<0.001). To further characterize the tdTO+ cells, we used different cell surface antibodies: CD31 (endothelial lineage), CD45 (hematopoietic lineage), Sca1 (mesoderm), VCAM1 (conventional Pax7+ MuSCs) to label these cells (Liu et al., 2015). Flow cytometry analysis revealed that tdTO cells were mainly CD45-negative (~90%) (Figure 2D) and VCAM1-positive (>99.99%), like Pax7+ MuSCs (Liu et al., 2015). However, it should be noted that Lin28a-tdTO+ cells make up only a tiny fraction of the total pool of VCAM1+ cells (0.73%). The difference is that conventional MuSCs are exclusively CD31- and Sca1-, whereas ~60% of Lin28a+ cells were positive for CD31 and ~70% were positive for Sca1, while the remainder were CD31- Sca1- like conventional Pax7+ MuSCs (Figure 2D). These results suggest that the majority of Lin28a+ cells might be more primitive VCAM1+ CD31+ Sca1+ mesodermal progenitors.

**Figure 2.**
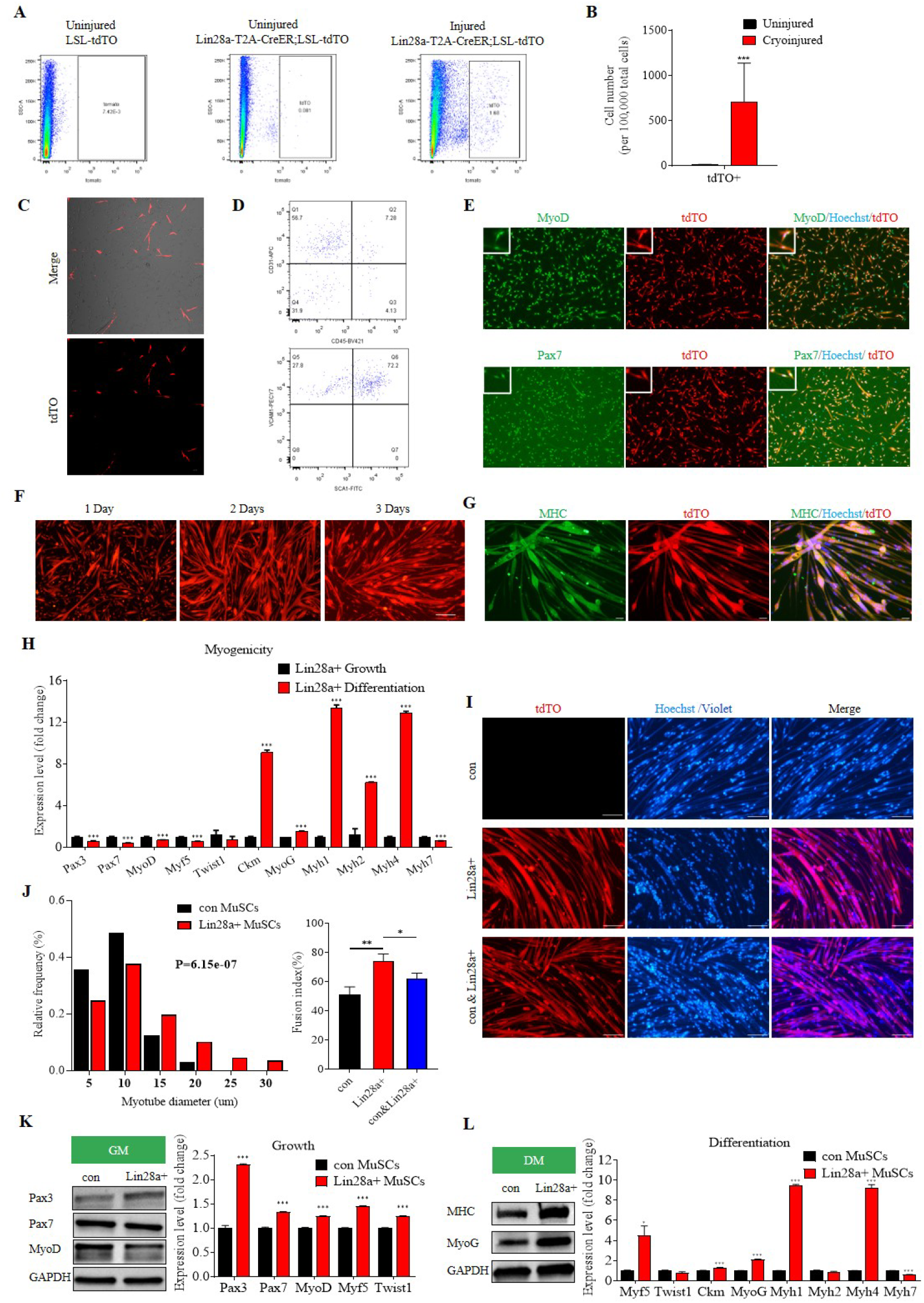
Lin28a+ cells are MuSCs which show enhanced myogenic potential in vitro. (**A**) Flow cytometry analysis for Lin28a-tdTO+ cells in the uninjured and injured muscles of Lin28a-T2A-CreER;LSL-tdTO mice. The control group was uninjured LSL-tdTO mice. All the mice were injected with TMX, and harvested 14 days after. (**B**) Quantification of the number of tdTO+ cells in injured or uninjured Lin28a-T2A-CreER;LSL-tdTO mice. N = 6 mice for each group. (**C**) Confocal microscopy on freshly FACS-isolated Lin28a-tdTO+ cells. Scale bar 50 μm. (**D**) Flow cytometry analysis for tdTO+ cells. Cells were first labeled with antibodies conjugated with fluorescent dyes for CD31 (APC), CD45 (BV421), VCAM1 (PE), Sca1 (FITC). The Lin28a-tdTO+ cells are mainly VCAM1+ CD31+ Sca1+ CD45- cells. (**E**) Pax7 and MyoD immunofluorescence staining (green) revealed that the majority of Lin28a+ cells cultured in growth medium (GM) expressed Pax7 and MyoD. Scale bar: 100μm. (**F**) Lin28a+ cells fused, differentiated and formed multinucleated myotubes starting at 1 and 2 days in differentiation medium (DM). Scale bar: 100μm. (**G**) Myosin Heavy Chain (MHC) (green) and Hoechst (blue) staining revealed that the majority of Lin28a-tdTO+ cells expressed MHC after differentiation into myotubes. Scale bar: 50μm. (**H**) Quantitative RT-PCR revealed that myogenic differentiation related genes, such as MyoG, Ckm, Myh1, Myh2, Myh4, were strongly activated, while myogenic progenitor-related genes, such as Pax3, Pax7, MyoD, Myf5, were significantly reduced, when Lin28a+ cells were cultured in differentiation media for 7 days, relative to undifferentiated Lin28a+ cells cultured in growth media. (**I**) Immunofluorescence images of conventional (con) MuSCs (CellTrace Violet-labeled), Lin28a+ cells, and a 1:1 mixture of con MuSCs with Lin28a+ cells, after FACS-isolation from Lin28a-T2A-CreER;LSL-tdTO mice’ TA muscles 14 days post-injury, then differentiated into myotubes for 36h. Scale bar: 100um. (**J**) Relative frequency distribution of the diameters of myotubes formed by the fusion of Lin28a+ cells and/or con MuSCs. For each bin, 200 myotubes were quantified, P =6.15 × 10^−7^. Right: Quantification of the fusion index of these myotubes. For each group, at least 3 different fields were quantified. (**K**) Western blot analysis for Pax3, Pax7, and MyoD protein expression in Lin28a+ cells, quantitative RT-PCR revealed expression levels of Pax3, Pax7, MyoD, Myf5 and Twist1 in Lin28a+ cells relative to con MuSCs, while proliferating in growth media (GM). Gapdh served as the loading control. (**L**) Western blot analysis for MHC and MyoG protein expression in Lin28a+ cell-derived myotubes, quantitative RT-PCR revealed expression levels of Myf5, Twist1, Ckm, MyoG, Myh1/2/4/7 in Lin28a+ cell-derived myotubes, relative to con MuSC-derived myotubes, while differentiating in DM. Gapdh served as the loading control.* P < 0.05, ** P < 0.01, *** P < 0.001.

Given these surface marker profiles, we attempted to differentiate freshly sorted Lin28a+ cells into different lineages in various differentiation media for skeletal myotubes, vascular endothelial cells, adipocytes or osteoblasts. The results showed that after Lin28a+ cells were differentiated in adipogenic medium, osteogenic medium, and endothelial cell medium, most of the cells underwent senescence or apoptosis, and could not further differentiate, although a fraction (~20%) did differentiate into alkaline phosphatase-positive osteoblasts in osteogenic medium (Figures S2B-E). In contrast, all the cells retained MyoD expression, suggesting that they maintained myogenic differentiation potential even in the presence of other lineage differentiation cues (Figures S2B-E).

In order to further compare the muscle differentiation potential of Lin28a+ cells with VCAM1+ CD31- Sca1- Pax7+ MuSCs (termed conventional MuSCs hereafter), we proliferated and differentiated Lin28a+ cells in myogenic growth media and differentiation media respectively, then performed immunofluorescence staining. The results showed that 100% of Lin28a+ cells could express the muscle stem/progenitor cell markers MyoD and Pax7 when proliferating (Figure 2E), and they could robustly form multinucleated myotubes expressing myosin heavy chain (MHC) proteins upon differentiation (Figures 2F-L). In the process of myogenic differentiation, the expression of many muscle stem/progenitor cell markers such as Pax3, Pax7, MyoD, Myf5 were downregulated, while the expression of many myogenic differentiation-related genes such as muscle creatine kinase (Ckm) and the myosin heavy chains (Myh1, Myh2 and Myh4) were upregulated significantly (Figure 2H). These results indicate that Lin28a+ cells proliferate as skeletal muscle progenitors in vitro, and are capable of myogenic fusion and differentiation into multi-nucleated myotubes.

In order to compare the fusion efficiency of Lin28a+ cells with conventional MuSCs, we cultured Lin28a+ cells, conventional MuSCs and 1:1 mixed cells, and measured their fusion index upon myogenic differentiation. Interestingly, we observed that Lin28a+ cell-derived myotubes were thicker in diameter and thus more hypertrophic, compared to conventional MuSC-derived myotubes (Figures 2I,J; P = 6.15 × 10^−7^). Furthermore, our results showed that Lin28a+ cells had a higher fusion index than conventional MuSCs, suggesting that Lin28a+ cells have higher myogenicity than conventional MuSCs (Figure 2J).

To further compare the molecular differences between Lin28a+ cells and conventional MuSCs, we compared the two groups’ myogenic factor expression by qRT-PCR and Western blot analysis. We found that, compared with conventional MuSCs, Lin28a+ cells expressed more Pax3 protein (Figure 2K), while the levels of Pax7 protein and MyoD protein were similar during proliferation (Figure 2K). Upon differentiation, Lin28a+ cell-derived myotubes expressed more MHC protein and MyoG protein, than conventional MuSC-derived myotubes (Figure 2L). Quantification of mRNA expression patterns reflected a similar pattern, where Lin28a+ cells expressed significantly higher levels of Pax3 than conventional MuSCs in growth media (Figure 2K). After terminal differentiation, Lin28a+ cell-derived myotubes expressed higher levels of Myf5, MyoG, Ckm, Myh1, and Myh4 than conventional MuSC-derived myotubes (Figure 2L). Notably, these pre-programmed differences in myogenicity persisted even after Lin28a expression is extinguished upon extended culture of Lin28a+ cells in vitro, as the expression of Lin28a protein and mRNA could not be detected in either growth media or differentiation media upon culture (Figures S3A, B). Taken together, these results indicate that Lin28a+ cells are distinct from conventional MuSCs in their pre-programmed myogenicity, at both the functional and molecular levels.

### Epigenomic and transcriptomic profiles show Lin28a dedifferentiates MuSCs

Lin28a and Pax3 are both typically expressed during embryogenesis and fetal development. Given that previous studies had suggested a DNA methylation epigenetic clock exists to monotonically mark biological aging from embryonic stem cells to adult cells to aged cells in mammals^24, 25^, we were motivated to understand the epigenetic profile of Lin28a+ cells at the genome-wide level. Thus we compared Lin28a+ MuSCs to embryonic and adult Pax7+ MuSCs (N = 3 mice each) through whole-genome bisulfite sequencing (WGBS; Figure S3D). Hierarchical clustering indicated that, while adult Lin28a+ MuSCs were more similar to adult Pax7+ MuSCs, they were also somewhat similar to embryonic Pax7+ myoblasts (Figure 3A). There were 5 clusters of differentially methylated regions (DMRs), instead of the 8 clusters expected by random chance, suggesting lower entropy and higher information structure amongst the MuSC subsets than expected (Figure 3A). DMR clusters 1 and 4 indicated the similarities between the two types of adult MuSCs. Cluster 3, consisting of 6573 DMRs and 5883 genes, indicated the surprising similarities between adult Lin28a+ MuSCs and embryonic Pax7+ myoblasts. DMR clusters 2 and 5 indicated the epigenetic uniqueness of the Lin28a+ cells as a distinct subset of MuSCs.

**Figure 3.**
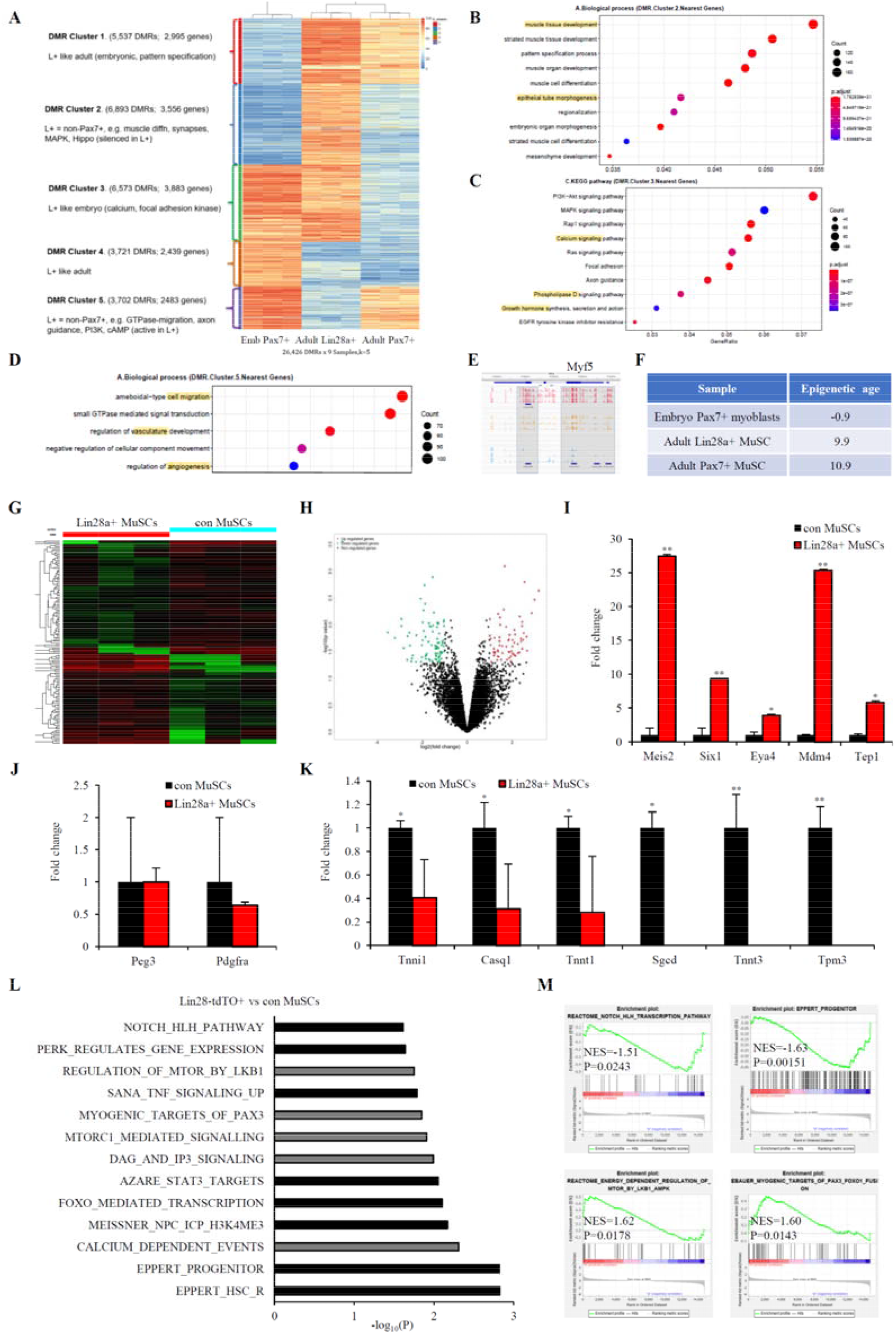
Epigenomic and transcriptomic profiles show Lin28a maintains MuSCs in a dedifferentiated state. (**A**) Hierarchical clustering analysis for the DNA methylomes of adult Lin28a+ MuSCs, adult Pax7+ MuSCs and embryonic (emb) Pax7+ myoblasts. N = 3 mice for each group. Differentially methylated region (DMR) clusters and summaries of Lin28a+ MuSC (L+) features are specified on the left. (**B**) GO analysis of Cluster 2 in (A). (**C**) KEGG analysis of Cluster 3 in (A). (**D**) GO analysis of Cluster 5 in (A). (**E**) CpG site methylation level in the region near the *Myf5* gene locus of Lin28a+ MuSCs (red), adult Pax7+ (yellow) and embryonic Pax7+ (blue) MuSCs (signal value 0-100%). Differentially methylated regions are highlighted in grey. (**F**) Application of a mouse multi-tissue epigenetic clock^29,30^ to our WGBS datasets, to predict adult Pax7+ and Lin28a+MuSCs’ and embryonic Pax7+ myoblasts’ epigenetic age. (**G**) Hierarchical clustering analysis for the transcriptomes of Lin28a+ MuSCs and conventional (con) MuSCs. N = 3 mice for each group. (**H**) Volcano plot analysis for differentially expressed genes in Lin28a+ MuSCs, relative to con MuSCs. Red, upregulated>2-fold and P<0.05. Green, downregulated>2-fold and P<0.05. (**I**) Expression levels of three primitive limb mesodermal progenitor transcription factors (Meis2, Six1, Eya4), Tep1 and Mdm4 in Lin28a+ MuSCs relative to con MuSCs. (**J**) Expression levels of markers expressed in other unconventional Pax7-independent muscle progenitors, Peg3 and Pdgfra, in Lin28a+ MuSCs relative to con MuSCs. P > 0.05. (**K**) Expression levels of myogenic terminal differentiation markers, including many troponins (Tnni1, Tnnt1, Tnnt3), calsequestrin (Casq1), sarcoglycan (Sgcd), and tropomyosin (Tpm3), in Lin28a+ MuSCs relative to con MuSCs. ** P<0.01, *P<0.05. (**L**) Signatures enriched in Lin28a+ MuSCs (black) or con MuSCs (grey), as identified by Gene Set Enrichment Analysis. **(M)** Representative GSEA profiles with normalized enrichment scores (NES) and nominal P values are shown. * P < 0.05, ** P < 0.01.

In particular, DMR cluster 2 was highly enriched for genes in muscle development and epithelialization according to GO analysis (Figures 3B and S3E), suggesting that Lin28a+ cells have silenced many genes for skeletal muscle differentiation and that prior to induction cues for terminal differentiation, Lin28a+ MuSCs are more dedifferentiated than Pax7+ MuSCs. Cluster 3 was highly enriched for genes involved in adult muscle-associated calcium signaling for contractility, axon guidance signaling for neuromuscular junctions, phospholipase D signaling^26^ and growth hormone signaling for hypertrophic growth, according to KEGG analysis (Figures 3C and S3F). These WGBS results confirmed that adult Lin28a+ MuSCs resembled embryonic Pax7+ myoblasts in silencing many adult muscle-associated genes. Cluster 5 was highly enriched for genes involved in cell migration and vascularization or angiogenesis (Figures 3D and S3G), suggesting that Lin28a+ cells resembled primitive somitic progenitors which are highly migratory and pro-angiogenic^27, 28^. However, the Lin28a+ MuSCs are still of adult origin like adult Pax7+ MuSCs, as exemplified by their similar methylation patterns of the myogenic transcription factor *Myf5* (Figure 3E). When we applied a mouse multi-tissue epigenetic clock model^29, 30^ to these WGBS datasets, we found that embryonic Pax7+ myoblasts were near-zero in their epigenetic age, while Lin28a+ MuSCs laid between embryonic Pax7+ myoblasts and adult Pax7+ MuSCs (Figure 3F). Overall, Lin28a+ MuSCs were more similar to adult Pax7+ MuSCs, but possessed unique and embryonic-like features that suggested dedifferentiation at the epigenomic level.

We then performed RNA-seq analysis to confirm the similarities and differences between adult Lin28a+ and Pax7+ MuSCs. Hierarchical clustering analysis (N = 3 mice each) showed that Lin28a+ cells differed widely from conventional Pax7+ MuSCs in their transcriptomes (Figure 3G). Volcano plot analysis showed that 78 genes were significantly upregulated >2-fold (P<0.05), while 83 genes were significantly downregulated >2-fold (P<0.05) (Figure 3H). Amongst the upregulated genes are 3 transcription factors that signify primitive limb bud mesoderm progenitors ^31–36^ : Meis2 (~27-fold), Six1 (~10-fold), and Eya4 (~3-fold), suggesting that Lin28a+ cells are similar to fetal limb muscle progenitors (Figure 3I). Furthermore, Lin28a+ cells also had higher Mdm4 (~25-fold) and Tep1 (~6-fold), suggesting more active telomerase and less active p53 activities (Figure 3I). Interestingly, prominent *let-7* targets such as Igf2bp2 and Hmga2 were not significantly upregulated (Figures S3H, I). We also examined the expression of markers expressed in other unconventional Pax7-independent muscle progenitors, Peg3 and Pdgfra^20, 37^, and found no significant increase compared to conventional Pax7+ MuSCs (Figure 3J). In contrast, markers of myogenic terminal differentiation were amongst the most significantly downregulated genes, including the troponins and tropomyosins (Figure 3K), confirming that Lin28a+ MuSCs are dedifferentiated relative to Pax7+ MuSCs.

Gene Set Enrichment Analysis (GSEA) showed that several stem cell signatures were upregulated in Lin28a+ cells, including the Notch pathway, the signature for neural progenitor cells (Meissner_NPC_ICP_H3K4me3), and the signatures for hematopoietic stem and progenitor cells (Eppert_HSC_R and Eppert_Progenitor) (P<0.05; Figure 3L). Many stress response pathways were also upregulated in Lin28a+ cells: the unfolded protein response mediated by PERK, the pro-inflammatory stress responses mediated by TNF-NFκB and IL6-STAT3, and the oxidative stress response mediated by FoxO (Figure 3L). In contrast, the mTOR pathway components, Ca^2+^ signaling-related signatures, and the myogenic differentiation markers (Myogenic_Targets_of_Pax3), all of which signify myogenic terminal differentiation, were downregulated in Lin28a+ cells compared to conventional MuSCs (Figures 3L,M). Our analysis results suggested that Lin28a+ cells show increased dedifferentiation and stress responsiveness, compared to conventional Pax7+ MuSCs.

To test if these signatures are just correlated with Lin28a expression or causally due to Lin28a expression, we overexpressed Lin28a in conventional MuSCs, and repeated the analysis (Figure S3C). GSEA showed that Lin28a overexpression led to increased stemness signatures again, such as the Wnt, Notch and Hedgehog signaling pathways, and the E2F-related mitosis or DNA replication signatures (Figures S3K,L). Interestingly, Lin28a also upregulated a hypoxia signature, i.e. genes that are downregulated upon HIF1A RNAi (Manalo_Hypoxia). In contrast, the myogenic differentiation signatures (Myogenic_Targets_of_Pax3; Striated_Muscle_Contraction) were significantly down-regulated by Lin28a (Figures S3K, L), indicating that Lin28a promotes stem cell self-renewal and dedifferentiation in MuSCs.

### Lin28a promotes the self-renewal of MuSCs and rejuvenates aged human muscle progenitors

Given that Lin28a expression is extinguished after Lin28a+ cells are cultured in vitro, we reactivated Lin28a expression in both Lin28a-tdTO+ cells and conventional MuSCs to explore the function of Lin28a in muscle stem cells. Compared with the empty vector control and conventional MuSCs, the proliferative self-renewal capacity of Lin28a-overexpressing tdTO+ cells were slightly stronger (Figures 4A, B). These findings were also confirmed by qRT-PCR (Figure 4C,D), which showed that after Lin28a overexpression, genes related to differentiation such as Ckm, MyoG, Myh1, Myh2, Myh4 decreased significantly (Figure 4C,D).

**Figure 4.**
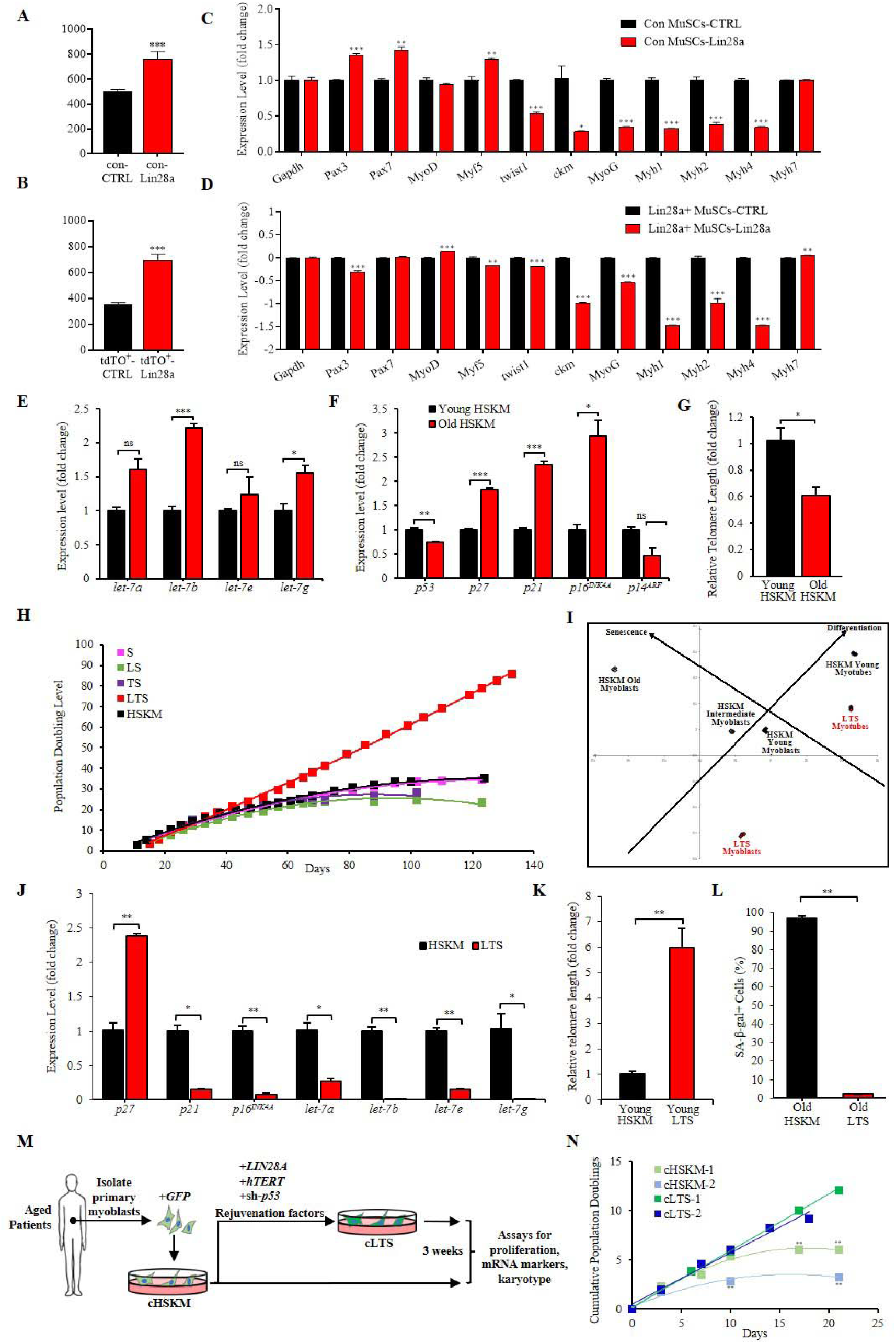
Lin28a promotes the self-renewal of MuSCs and aged human muscle progenitors. (**A-B**) Cell proliferation rate of con MuSCs (A) and Lin28a-tdTO+MuSCs (**B**), after being infected with retroviruses expressing either empty vector (CTRL) or Lin28a at P20 (con) or P10 (Lin28a+). (P indicates passages). (**C-D**) Quantitative RT-PCR for myogenic differentiation markers in con MuSCs (C) or Lin28a-tdTO+ MuSCs (D) which overexpressed Lin28a, relative to con MuSCs or Lin28a-tdTO+ MuSCs with the empty vector (CTRL). Data are mean ± SEM. N = 3 independent experiments. (**E**) Quantitative RT-PCR for *let-7* microRNAs in aged adult HSKM myoblasts, relative to young adult HSKM myoblasts. (**F**) Quantitative RT-PCR for mRNAs of cell cycle inhibitors in aged adult HSKM myoblasts, relative to young adult HSKM myoblasts. (**G**) Quantitative RT-PCR for telomere length in aged adult HSKM myoblasts, relative to young adult HSKM myoblasts. (**H**) Population doubling curves for young HSKM myoblasts (black), and young adult HSKM myoblasts transduced with viral sh-p53 (S, pink), and Lin28a (LS, green), or hTERT (TS, purple), or Lin28a and hTERT (LTS, red). While young adult HSKM and other transgenic myoblasts started to undergo senescence by 60 days, before the 30^th^ population doubling, the LTS myoblasts (red) continued to proliferate steadily beyond 120 days and beyond the 90^th^ population doubling. (**I**) Principal Components Analysis (PCA) of transcriptomic data for LTS myoblasts and myotubes (red), relative to HSKM young, intermediate and old myoblasts, and young HSKM myotubes (black). (**J**) Quantitative RT-PCR for *p27^KIP1^, p21^CIP1^, p16^INK4A^*, and *let-7* microRNAs in LTS myoblasts, relative to young adult HSKM myoblasts. (**K**) Quantitative RT-PCR for telomere length in young LTS myoblasts, relative to young adult HSKM myoblasts. (**L**) Quantification of senescence-associated β-galactosidase positive (SA-β-gal+) cells in 100-day-old adult HSKM myoblasts, relative to 100-day-old LTS myoblasts. (**M**) Schematic for derivation and rejuvenation of aged patients’ myoblasts. (**N**) Population doubling curves for two representative aged patients’ HSKM myoblasts (cHSKM-1, cHSKM-2), and their corresponding LTS-treated lines (cLTS-1, cLTS-2). Ns, not significant, * P < 0.05, ** P < 0.01, *** P < 0.001.

Furthermore, we also tested whether the lineage potential of tdTO+ cells was altered by Lin28a overexpression, by exposing the cells to media for differentiation towards adipocytes, osteoblasts, and endothelial cells. According to qRT-PCR results, we found that the overexpression of Lin28a enhanced the capacity for osteogenic differentiation, weakly but significantly inhibited the capacity for adipogenic differentiation, and produced little effect on vascular cell differentiation (Figure S3J). These results suggest that the reactivation of Lin28a in cultured tdTO+ cells can partially mimic the characteristics of freshly sorted Lin28a+ cells in vivo, which also showed a weak propensity for osteogenesis but not adipogenesis nor vascular differentiation (Figures S2 and S3J). Taken together, these results suggest that the function of Lin28a in Lin28a+ MuSCs may be to promote self-renewal while retaining enhanced myogenic and even osteogenic potential.

Given that aged human myoblasts lose their self-renewal and myogenic potential rapidly during senescence, that human myoblasts accumulate the Lin28-targeting *let-7* microRNAs during development and aging (Figure 4E) ^39^, and to test the human relevance of our mouse lineage-tracing findings, we attempted to overexpress Lin28a in adult human skeletal muscle (HSKM) myoblasts as well. First, we found that old adult HSKM myoblasts, compared to young myoblasts, showed significantly higher levels of cell cycle inhibitors such as *p21^CIP1^*, *p27^KIP1^*, and *p16^INK4a^* (Figure 4F), as well as p53 protein (Figure S4A). In contrast, the telomeres of old HSKM myoblasts were significantly shorter than young HSKM myoblasts (Figure 4G). Interestingly, we had found that the P53 inhibitor Mdm4 and the telomerase complex component Tep1 were amongst the most highly upregulated genes in Lin28a+ MuSCs as well (Figure 3C). These results spurred us to biomimic Lin28a+ MuSCs by conducting a mini-screen of factors along with Lin28a to attempt to rejuvenate and/or dedifferentiate human adult myoblasts.

Human adult myoblasts eventually underwent replicative senescence, plateauing sigmoidally by ~30 population doublings, and no single factor alone could prevent the senescence (Figures 4H and S4B). However, when Lin28a was combined with hTERT and a short hairpin against *TP53* (LTS; Figure S4C), human adult myoblasts could self-renew and proliferate indefinitely, beyond 90 population doublings (currently over 200 doublings), with a linear doubling curve throughout. Furthermore, each factor was necessary in the LTS combo, as omission of any one of the three factors resulted in either senescence or massive apoptosis (e.g. Lin28a alone or hTERT alone or both), thus causing the population doubling curve to plateau early (Figure 4H).

We compared LTS to normal primary HSKM cells, both myoblasts and myotubes, and both young and old, using genome-wide transcriptomic profiling to assess their cellular states. Principal components analysis (PCA) of the multi-dimensional mRNA space revealed that the different cell-types were clearly separated by their senescence and differentiation states (Figure 4I). The differentiation vector, defined between HSKM myoblasts and myotubes, was approximately parallel to the LTS cells’ differentiation vector, proving again that the myogenesis program was conserved after LTS treatment. The senescence vector defined between the HSKM young, intermediate, and old myoblasts, was approximately orthogonal to the differentiation vector, proving that terminal differentiation and senescence are two related but distinct concepts, even if both processes involved a cell cycle exit. Based on the differentiation and senescence vectors defined by the primary HSKM cells, LTS myoblasts appeared rejuvenated and dedifferentiated, compared to young adult primary myoblasts (Figure 4I).

When examined in detail, we found that the LTS factors prevented the pro-senescent induction of *p21^CIP1^*, *p16^INK4a^*, *let-7* microRNAs (Figure 4J), and P53 protein (Figure S4A), although *p27^KIP1^* remained high. The LTS combo also lengthened the telomeres significantly (Figure 4K).

When 100 day-old adult HSKM myoblasts and LTS myoblasts were examined for senescence-associated β-galactosidase (SA-β-gal) staining, 96.7% of normal adult progenitors were SA-β-gal+, whereas only 2.2% of LTS progenitors were SA-β-gal+, indicating that the LTS combo was profoundly protective against senescence (Figures 4L, S4D).

To test the utility of the LTS factors in aged muscle progenitors, we transduced muscle progenitors from aged patients with the LTS combo (Figure 4M). When we cultured aged patients’ HSKM myoblasts, we found that they rapidly underwent senescence within 6 population doublings (Figure 4N), which is significantly less than healthy young adult HSKM progenitors (~30 population doublings, Figure 4H). However, upon treatment with the LTS factors (Figure S4E), the aged HSKM progenitors successfully averted rapid senescence with linear population doubling curves (Figure 4N). By now, we have generated over 20 patient myoblast cell-lines that can grow past 160 population doublings.

### Lin28a promotes MuSC dedifferentiation and muscle regeneration in vivo

We characterized the differentiation capacity of LTS progenitors under serum withdrawal conditions, to test if they had undergone oncogenic transformation in vitro. In growth conditions, LTS progenitors reactivated Pax3 protein (Figure 5A), showed higher expression of MyoD (Figure 5B), lower expression of many myogenic differentiation markers (Figures 5C-F, S5A), and appeared dedifferentiated overall, similar to the Lin28a+ cells in vivo (Figures 3, S3J). But unlike immortalized progenitors^40^ harboring oncogenic CDK4^R24C^ and 100-day-old senescent HSKM progenitors, which both failed to differentiate properly after serum withdrawal (Figure S4F), 100-day-old LTS progenitors could still undergo terminal differentiation into myosin heavy chain (MHC)+, α-actinin+, myogenin+ multi-nucleated myotubes like young adult HSKM progenitors (Figures S4F, G).

**Figure 5.**
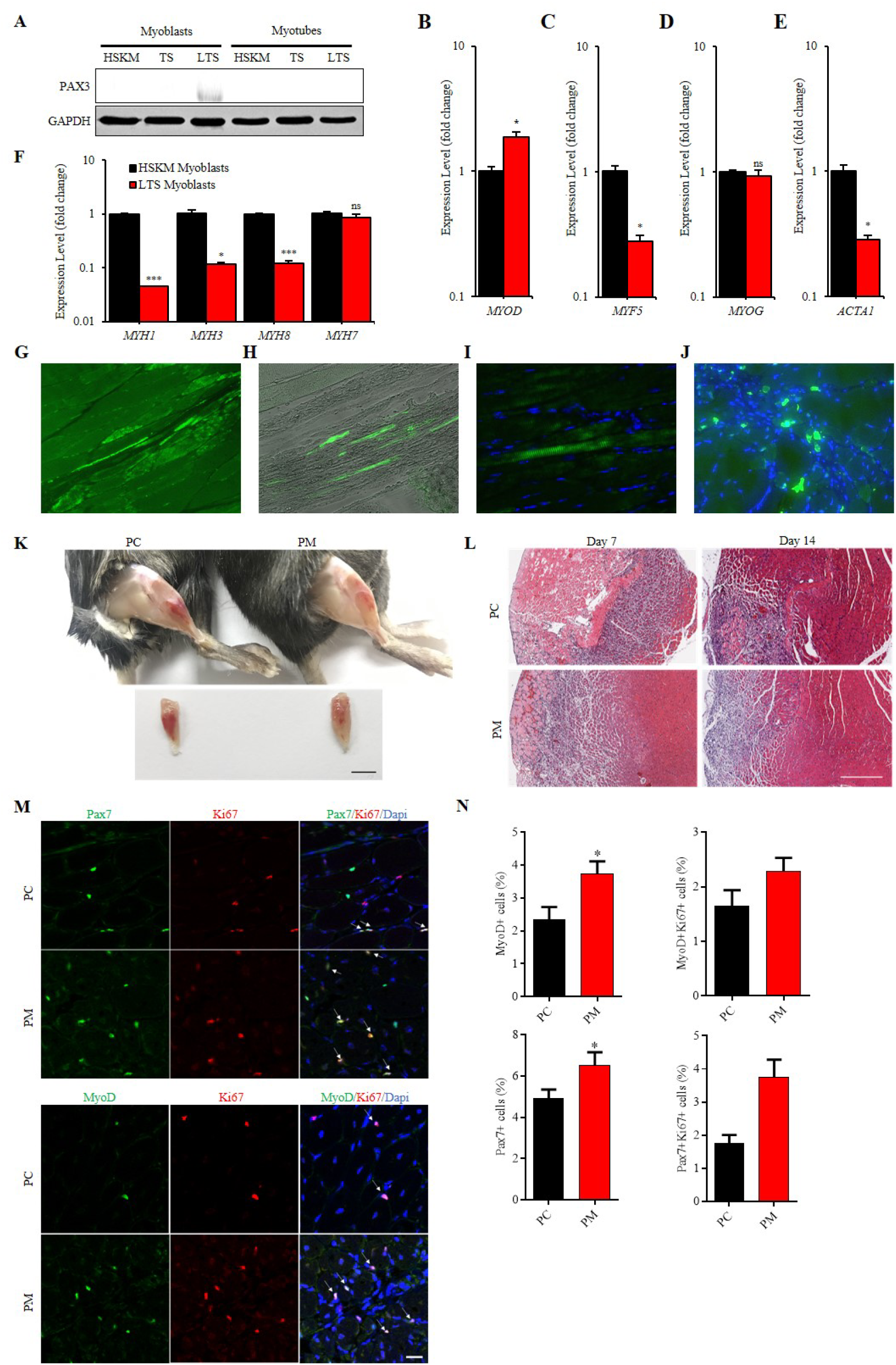
Lin28a promotes MuSC dedifferentiation and muscle regeneration in vivo. **(A)** Western blot for PAX3 protein, relative to GAPDH protein, in young HSKM, TS and LTS myoblasts and myotubes. **(B)** Quantitative RT-PCR for *MYOD1* in LTS myoblasts, relative to young adult HSKM myoblasts. **(C)** Quantitative RT-PCR for *MYF5* in LTS myoblasts, relative to young adult HSKM myoblasts. **(D)** Quantitative RT-PCR for the differentiation marker *MYOG* in LTS myoblasts, relative to young adult HSKM myoblasts. **(E)** Quantitative RT-PCR for the differentiation marker *ACTA1* in LTS myoblasts, relative to young adult HSKM myoblasts. **(F)** Quantitative RT-PCR for the terminal differentiation markers *MYH1, MYH3, MYH8, MYH7* in LTS myoblasts, relative to young adult HSKM myoblasts. (**G**-**J**) Immunofluorescence staining for GFP showed that there was significant engraftment of LTS-treated aged progenitors, 16 days after orthotopic injection into the gastrocnemius muscles of NSG mice after cryoinjury. (**G, H**). GFP+ LTS human progenitors had mostly differentiated into elongated myofibers, or (**I**) some fused with mouse myofibers to form GFP+ domains within the myofibers, and **(J)** some persisted as mononuclear self-renewing progenitors on the periphery of the myofibers. **(K)** Photos show the resolution of muscle inflammation and muscle regeneration in the TA muscles of Pax7-CreERT2 (PC) and Pax7-CreER; NFKB-LSL-Lin28a (PM) mice on day 7 post injury. Scale bar: 5mm. **(L)** Hematoxylin and eosin (H&E) staining of TA muscle sections of PM mice, relative to PC mice, on days 7 and 14 after cryoinjury. Scale bar 300μm. **(M)** Immunofluorescence staining for MyoD, Pax7 and Ki67 expression in the damaged TA muscles from PM mice, relative to Pax7-CreERT2 mice. Scale bar 20μm. **(N)** Quantification of the percentage of myoblasts, after injured TA muscle sections were stained for MyoD and Pax7. For each muscle section, at least 7 different fields were quantified. Ns, not significant, * P < 0.05, ** P < 0.01, *** P < 0.001.

We were also interested to test if the LTS factors would lead to genomic instability, especially in aged progenitors, given the inhibition of *TP53*. When we subjected late-passage LTS progenitors to chromosomal analysis, the cells still displayed a normal diploid karyotype (Figure S5B). In contrast, late-passage TS progenitors displayed a variety of aneuploid karyotypes, including the loss of numerous chromosomes, and the appearance of dicentric, ring and marker chromosomes (Figure S5C). These results suggested that Lin28a might promote chromosomal stability even after *TP53* inhibition in muscle progenitors, and not tumorigenicity.

To functionally test if LTS progenitors can also engraft to contribute to muscle regeneration like MuSCs, instead of tumorigenesis in vivo, GFP+ LTS progenitors were massively expanded then injected orthotopically into the gastrocnemius muscles of immunodeficient mice after cryoinjury. Immunofluorescence staining for GFP showed that there was significant engraftment of LTS progenitors 30 days after injection (Figures 5G-J). And the GFP+ LTS human progenitors had mostly differentiated into elongated myofibers (Figures 5G-H), or fused with mouse myofibers to form GFP+ domains (Figure 5I), and some persisted as self-renewing muscle progenitors on the periphery of the myofibers (Figure 5J). Thus, LTS progenitors can contribute to muscle regeneration in vivo, without forming tumors.

Our in vitro experiments have shown that Lin28a overexpression promotes the self-renewal and dedifferentiation of MuSCs without compromising their myogenic capacity, which led us to ask if overexpression of an injury-responsive Lin28a cassette in adult Pax7+ muscle stem cells can also enhance muscle regeneration in vivo. Therefore, we constructed the NFKB-loxp-stop-loxp-Lin28a transgenic mouse (no. mt190), and bred it with Pax7-CreERT2 mice to obtain heterozygous PM (Pax7-CreERT2; mt190) mice that can express Lin28a in Pax7+ MuSCs only during muscle injury, and shut it off after inflammatory signaling resolves.Following the same schedule of injection of TMX and injury as our lineage-tracing experiments (Figure 1A), we tested the regeneration ability of PM mice at 7 and 14 days after double-blinded cryoinjury tests on TA muscles. By 7 days after injury, it was already visually obvious that the resolution of reddish, necrotic and inflammed muscle tissue was more improved in PM mice than PC (Pax7-CreERT2) mice (Figure 5K).Hematoxylin and eosin staining of TA muscle sections revealed that PM mice’ TA muscles have smaller pink necrotic zones and larger regenerative zones with centrally nucleated myofibers, between 7 to 14 days after injury (Figure 5L). These results indicated that the muscle regeneration capacity of PM mice was significantly enhanced by Lin28a.

In order to determine which phase of muscle regeneration was enhanced in PM mice, we performed immunofluorescence staining on the TA muscle sections. We found that, consistent with our observations that Lin28a overexpression rejuvenated MuSCs, the numbers of proliferating Pax7+ MuSCs and MyoD+ myoblasts were significantly higher in the TA muscles of PM mice than in PC mice (Figures 5N, O). These results suggest that the injury-induced overexpression of Lin28a in Pax7+ MuSCs promotes their self-renewal and proliferation, thereby enhancing skeletal muscle regeneration capacity in response to injuries.

### Lin28a optimizes OxPhos and mtROS to induce HIF1A-glycolysis

To dissect the mechanism for Lin28a’s extension of muscle progenitor self-renewal capacity, we used adult primary HSKM progenitors cultured ex vivo. Our RNA profiling studies earlier had shown that *let-7* microRNAs accumulate during ageing ^41^. Both TS and LTS transduction lowered the *let-7* levels, with Lin28a further suppressing *let-7b, let-7e*, and *let-7g* levels on a log scale (Figure S6A). To test if Lin28a’s promotion of self-renewal in muscle progenitors was mediated by *let-7*, we transfected mature let-7 mimics in an attempt to abrogate Lin28a’s dramatic effect on proliferation (Figure 6A). Western blots showed that important *let-7* targets that were derepressed by Lin28a ^41^, such as IGF2BP1/2/3 and HMGA2, were almost completely depleted by the *let-7* overexpression in human muscle progenitors (Figure S6B). However, we found that neither *let-7a* nor *let-7b*, nor both, had any effect on LTS progenitors’ proliferation rate (Figure 6A). Moreover, *let-7* overexpression also failed to increase the numbers of senescent SA-β-gal+ cells (Figure S6C). These results suggest that *let-7* repression alone cannot fully explain Lin28a’s effect on muscle progenitor self-renewal.

**Figure 6.**
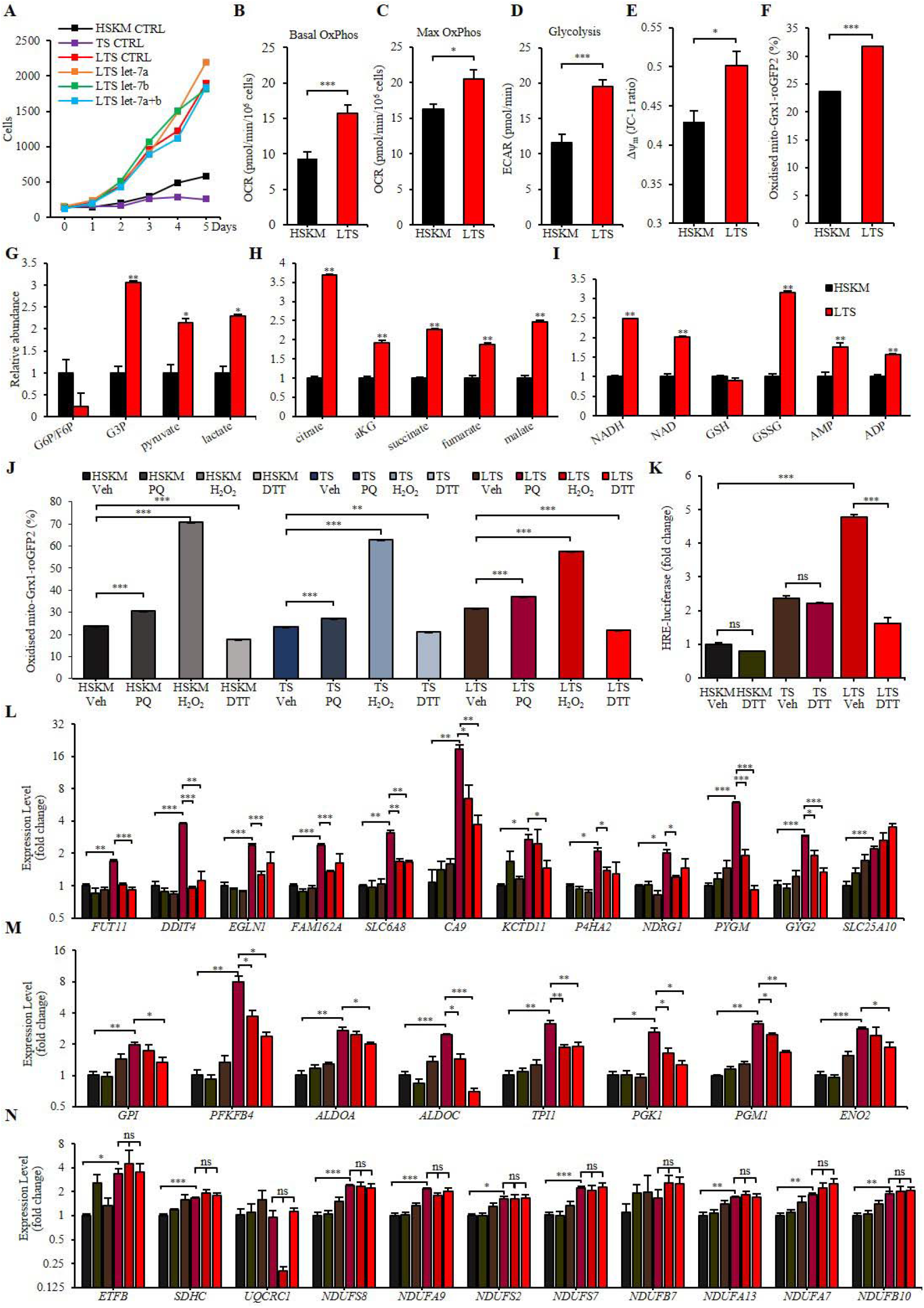
Lin28a optimizes OxPhos and mtROS to induce HIF1A-glycolysis. **(A)** Cell counts after control RNAi (CTRL), *let-7*a, *let-7*b, or combined *let-7*a and *let-7*b (*let-7*a+b) mimic transfection in TS and LTS myoblasts, relative to young adult HSKM myoblasts. **(B)** Seahorse analysis of maximal OxPhos in LTS myoblasts, relative to young adult HSKM myoblasts. **(C)** Seahorse analysis of glycolysis in LTS myoblasts, relative to young adult HSKM myoblasts. **(D)** Quantification of the mitochondrial membrane potential (Δψ_m_) in LTS myoblasts, relative to young adult HSKM myoblasts, according to the JC-1 dye red : green fluorescence ratio. **(E)** Quantification of mitochondrial ROS (mtROS) in LTS myoblasts, relative to young adult HSKM myoblasts, according to the mean percentage oxidation of the mito-Grx1-roGFP2 probe. **(F)** LC-MS/MS quantification of glycolysis metabolites in LTS myoblasts (red) and adult HSKM myoblasts (black). G6P, glucose-6-phosphate. F6P, fructose-6-phosphate. G3P, glyceraldehyde-3-phosphate. **(G)** LC-MS/MS quantification of Krebs Cycle metabolites in LTS myoblasts (red) and adult HSKM myoblasts (black). aKG, α-ketoglutarate. **(H)** LC-MS/MS quantification of redox and bioenergetics-related metabolites in LTS myoblasts (red) and adult HSKM myoblasts (black). NADH, reduced NAD. NAD, nicotinamide adenine dinucleotide. GSH, glutathione. GSSG, glutathione disulfide. AMP, adenine monophosphate. ADP, adenine diphosphate. **(I)** Gene Set Enrichment Analysis (GSEA) of transcriptomic profiles of LTS myoblasts. **(J)** Quantification of mitochondrial ROS (mtROS) in young adult HSKM, TS and LTS myoblasts, after treatment with the ROS-modulating drugs paraquat (PQ), hydrogen peroxide (H_2_O_2_), dithiothreitol (DTT), or vehicle (Veh), according to the mean percentage oxidation of the mito-Grx1-roGFP2 probe. **(K)** Quantification of the hypoxia-response element (HRE)-luciferase reporter in young adult HSKM, TS and LTS myoblasts, after treatment with dithiothreitol (DTT) or vehicle control. **(L)** Quantitative RT-PCR for hypoxia/HIF1A target mRNAs in LTS myoblasts, relative to TS myoblasts, after treatment with vehicle control (CTRL), hydrogen peroxide (H_2_O_2_), or dithiothreitol (DTT). **(M)** Quantitative RT-PCR for glycolysis-related mRNAs in LTS myoblasts, relative to TS myoblasts, after treatment with vehicle control (CTRL), hydrogen peroxide (H_2_O_2_), or dithiothreitol (DTT). **(N)** Quantitative RT-PCR for OxPhos-related mRNAs in LTS myoblasts, relative to TS myoblasts, after treatment with vehicle control (CTRL), hydrogen peroxide (H_2_O_2_), or dithiothreitol (DTT). Ns, not significant, * P < 0.05, ** P < 0.01, *** P < 0.001.

Besides regulating the *let-7* microRNAs, Lin28a also binds and regulates mRNAs^8,11,15^. Thus, we analyzed the transcriptomic profiles of early-passage LTS progenitors, and compared them against early-passage TS progenitors by GSEA. Curiously, the top downregulated signatures in LTS progenitors were almost completely dominated by interferon response targets (Figure S6D). This is consistent with findings that the interferon response is activated during senescence^42^, and suggests that TS progenitors were activating inflammaging-induced senescence, which Lin28a downregulated. Our analysis further revealed that Lin28a’s effect was primarily metabolic in nature, with the top upregulated signatures consisting of mitochondrial oxidative phosphorylation (OxPhos) genes, stress-responsive signatures such as the unfolded protein response (UPR) mediated by XBP1s activity, the DNA damage response mediated by p53 activity, the hypoxic stress response mediated by HIF1A and the glycolysis genes (Figure S6D).

To validate that OxPhos and glycolysis were indeed upregulated, we used the Seahorse extracellular flux analyzer. Our analysis of oxygen consumption rates in the LTS progenitors revealed that LTS progenitors do indeed show significantly higher basal and maximal OxPhos rates (Figures 6B, C). Moreover, LTS progenitors also showed higher glycolysis rates (Figure 6D). We also found that the mitochondrial membrane potential Δψ_m_ in LTS progenitors was significantly increased (Figure 6E). In contrast, mitochondrial DNA and protein biogenesis did not increase, indicating that while the mitochondrial OxPhos activity levels were increased, total mitochondrial biogenesis did not increase (Figures S6E, F).

Since Lin28a has already been shown to bind OxPhos mRNAs to regulate OxPhos protein expression ^11, 43, 44^, it is plausible that Lin28a-induced OxPhos would increase mtROS via an increased ETC flux and increased Δψ_m_. Previous studies had shown that a high mitochondrial membrane potential Δψ_m_ can shift the mitochondrial ROS-producing sites to a more reduced state, thereby increasing the propensity for mitochondrial ROS (mtROS) production ^45, 46^. To validate this hypothesis, we used the mitochondrial Grx1-roGFP2 reporter to determine mtROS levels in live cells ^47^. Our results showed that LTS progenitors had mildly but significantly higher mtROS than HSKM progenitors (Figure 6F). When we performed LC-MS/MS metabolomics profiling, we confirmed that several glycolytic intermediates, mitochondrial Krebs Cycle intermediates, and bioenergetic molecules were indeed increased in LTS progenitors (Figures 6G-I). LC-MS/MS metabolomics also showed that the oxidized to reduced glutathione (GSSG/GSH) ratio, an indicator of oxidative stress, was higher in LTS progenitors (Figure 6I).

Since mtROS is an activator of HIF1A, which in turn transactivates glycolysis genes, mtROS could explain why the hypoxia and glycolysis signatures were both upregulated ^46^. If Lin28a was indeed optimizing mtROS to prevent senescence, then ROS-modulating drugs should have a disproportionately bigger effect on LTS progenitors than TS or HSKM progenitors. To test this hypothesis, we subjected the various muscle progenitors to low doses of a variety of ROS-modulating drugs and examined their effects on mtROS, gene expression, and proliferation rates. First, we found that H_2_O_2_ most reliably increased mtROS, whereas DTT most reliably reduced mtROS in all progenitor types (Figure 6J). Using a HIF1A-response-element luciferase reporter ^48^, we validated that Lin28a does induce a HIF1A-mediated hypoxic response in LTS progenitors relative to TS and HSKM progenitors, and that this stress response was dependent on ROS levels (Figure 6K).

When we subjected the progenitors to H_2_O_2_ and DTT treatment, we found that both higher ROS and lower ROS could suppress the HIF1A targets (Figure 6L) and glycolysis genes (Figure 6M) in LTS progenitors, but not in TS progenitors. This indicates that Lin28a was driving an optimal mtROS level to upregulate HIF1A targets and glycolysis genes in LTS progenitors. In contrast, both H_2_O_2_ and DTT treatments failed to perturb the mildly higher OxPhos mRNA levels in LTS progenitors (Figure 6N), indicating that OxPhos mRNA regulation was upstream of the mtROS levels. These results support a model whereby Lin28a ➔ optimal OxPhos and mtROS ➔ HIF1A-driven hypoxic and glycolysis responses ➔ self-renewal. However, it remained unclear which of these effects are actually responsible for Lin28a’s effects on muscle progenitor self-renewal.

### Lin28a-mtROS-HIF1A promotes self-renewal

When we examined their proliferative capacity, we found that H_2_O_2_ and DTT treatments both led to significant suppression of LTS progenitor proliferation in a dose-dependent fashion, with no significant effects on TS or HSKM progenitor proliferation, supporting the notion that Lin28a promotes optimal mtROS to increase the proliferative capacity (Figure 7A). Consistent with the model above, we also found that treatment with the HIF1A inhibitor KC7F2 significantly suppressed LTS progenitor proliferation, with no significant effects on TS or normal HSKM progenitor proliferation (Figure 7B).

**Figure 7.**
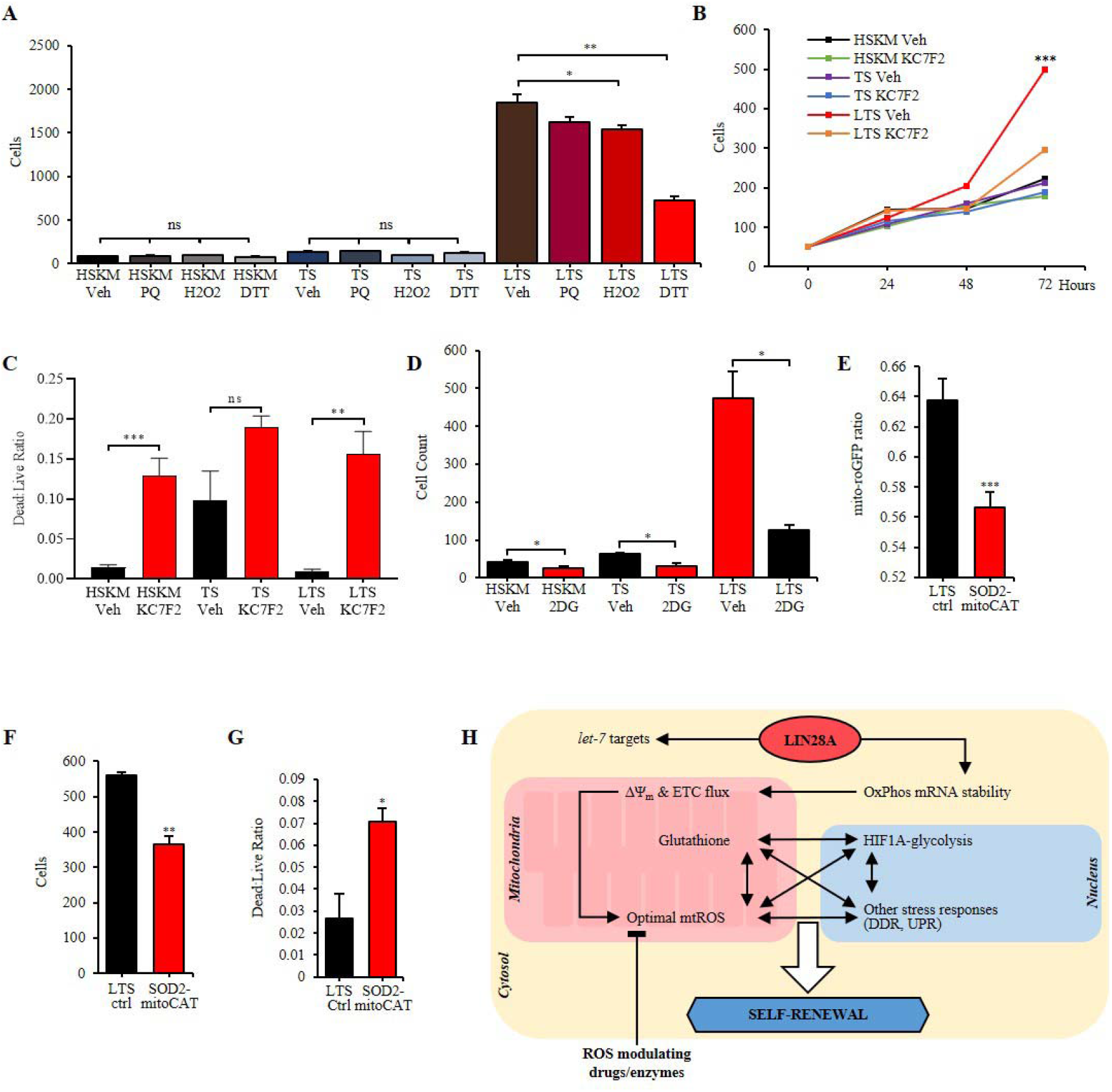
Lin28a-mtROS-HIF1A promotes self-renewal. **(A)** Cell counts after treatment with vehicle control (Veh), hydrogen peroxide (H_2_O_2_), or dithiothreitol (DTT) titrations in TS and LTS myoblasts, relative to young adult HSKM myoblasts. **(B)** Cell counts after treatment with vehicle control (Veh) or the HIF1A inhibitor KC7F2 in TS and LTS myoblasts, relative to young adult HSKM myoblasts. **(C)** Ratio of dead to live cells, as measured by ethidium and calcein staining, after treatment with vehicle control (Veh) or the HIF1A inhibitor KC7F2 in TS and LTS myoblasts, relative to young adult HSKM myoblasts. **(D)** Cell counts after treatment with vehicle control or the glycolysis inhibitor 2-deoxyglycose (2DG) in TS and LTS myoblasts, relative to young adult HSKM myoblasts. **(E)** Quantification of mitochondrial ROS (mtROS) in LTS myoblasts, before and after overexpression of mitochondria-targeted superoxide dismutase 2 and mitochondria-targeted catalase (SOD2-mitoCAT), according to the mito-Grx1-roGFP2 probe. **(F)** Cell counts before and after overexpression of SOD2-mitoCAT in LTS myoblasts. **(G)** Ratio of dead to live cells before and after overexpression of SOD2-mitoCAT in LTS myoblasts. **(H)** Model of how Lin28a, mtROS and the HIF1A-associated metabolic stress response network promotes stem cell self-renewal. Ns, not significant,* P < 0.05, ** P < 0.01, *** P < 0.001.

Live-dead staining of the cells revealed that in control vehicle-treated cells, both normal HSKM and LTS progenitors had less cell death than TS progenitors (Figure 7C), indicating that TS increased both mitosis and cell death, but Lin28a suppressed the cell death. Adding the HIF1A inhibitor KC7F2 increased the proportion of dead cells similarly across HSKM, TS and LTS progenitors (Figure 7C), suggesting that HIF1A was a common requirement for survival in all the muscle progenitors. Moreover, Lin28a-mediated suppression of cell death was over-ridden by KC7F2, suggesting that Lin28a promotes cell survival via HIF1A. The glycolysis inhibitor 2-deoxyglucose led to mild but significant decreases in all progenitors’ proliferation, but the decrease was most pronounced in LTS progenitors (Figure 7D), supporting the notion that Lin28a is driving HIF1A and glycolysis to promote proliferation.

To further confirm the role of mtROS in this mechanistic model, we used genetic perturbation of mtROS-modulating enzymes to examine the necessity of mtROS to Lin28a’s mechanism of action in cell proliferation. Targeted overexpression of mitochondrial superoxide dismutase 2 (SOD2) and mitochondrial catalase (mitoCAT) led to a significant decrease in mtROS (Figure 7E). This decrease in mtROS reduced the proliferation of LTS progenitors (Figure 7F), and also increased cell death (Figure 7G).

Taken together, these results demonstrate that Lin28a specifically optimized mtROS production and various stress responses (Figure 7H) to promote the self-renewal capacity and rejuvenation of adult muscle progenitors. Indeed, we observed lower levels of the DNA oxidative damage biomarker 8-oxoguanine in LTS progenitors (Figures S6G, H), as an effect of these stress responses (Figure S6D). These results are consistent with the stress-responsive signatures that we uncovered in mouse Lin28a+ MuSCs in vivo (Figure 3F), and suggests that the mitohormetic pathway is licensed to maintain a subset of MuSCs in a rejuvenated and dedifferentiated state in vivo.

## Discussion

Lin28a is mainly expressed during embryonic development and declines with development, while its ectopic overexpression can promote the regeneration of various adult tissues^11,41,49^. However, its expression and role(s) in endogenous adult tissue regeneration and dedifferentiation had remained elusive hitherto. Through lineage tracing, we have identified a group of previously unidentified MuSCs which express Lin28a and can respond to muscle damage with enhanced regenerative potential. Interestingly, this subset of satellite cells are not conventional Pax7+ MuSCs, as they possess epigenomic and transcriptomic profiles that suggest they lie between adult and embryonic Pax7+ muscle stem cells. In addition, we confirmed that Lin28a+ cells expressed more Pax3 and embryonic limb bud mesoderm transcription factors such as Meis2, Six1 and Eya4, compared with conventional Pax7+ MuSCs. Pax3 regulates limb muscle development by regulating Six1 and the Eya family^27, 35, 36, 50–53^, whereas Lin28a is expressed in the early stage of limb bud development during mouse embryogenesis^54, 55^. Consistent with these reports, we also traced Lin28a+ cells during embryonic limb bud development (Figure S7), and found migrating limb muscle progenitor cells that gave rise to limb bud myofibers. Lin28a upregulated Notch signaling for enhanced self-renewal and increased stress-responsive capabilities in the MuSCs, while suppressing terminal differentiation, further supporting the notion that Lin28a can reverse the epigenetic clock and stably maintain a dedifferentiated state. This extends our knowledge on Lin28a, which had been previously thought to merely promote cell proliferation during reprogramming and regeneration^5, 56, 57^.

Given their rarity (~0.7% of VCAM1+ cells), it is highly possible that the ablation of dedifferentiated Lin28a+ MuSCs may have no effect on steady-state adult muscle quality, quantity and hypertrophy. Although previous reports have found that Lin28a itself is not necessary for myogenesis^41^, it does not preclude the importance of Lin28a in maintaining a stably dedifferentiated state in a small number of MuSCs, as we have shown by lineage-tracing. Furthermore, previous findings have suggested that Lin28a is physiologically important in mouse limb bud and tail bud mesoderm development^55, 58^. In addition, we confirmed that injury-activated, NF-κB promoter-driven Lin28a expanded the numbers of Pax7+ and MyoD+ muscle progenitors, thereby accelerating necrosis resolution and improving muscle regeneration. Given our findings on Lin28a in human muscle progenitors, future work could focus on the translational value of Lin28a in rejuvenating muscle progenitors ex vivo.

In fact, we found that Lin28a not only promotes dedifferentiation and proliferation, but also the myogenic efficiency of MuSCs, which is important given that the myogenicity of MuSCs declines steadily with development and aging^59^. Nowhere is this decline more obvious than in old human muscle progenitors during human aging and sarcopenia^60^. We further showed that pre-senescent but old human muscle progenitors accumulate *let-7* microRNAs, *TP53*, *p21^WAF1^*, *p16^INK4a^*, and show telomere loss during aging. Given that we also found high levels of the p53 functional inhibitor Mdm4 and the telomerase complex component Tep1 in Lin28a+ MuSCs^61, 62^, we sought to biomimic Lin28a+ MuSCs in a mini-screen to rejuvenate old human muscle progenitors. Our screen revealed that a minimal set of Lin28a, telomerase and a shRNA against *TP53* (LTS) was sufficient to extend the self-renewal capacity of adult human muscle progenitors indefinitely. Given that telomerase and p53 inhibition are already used to immortalize cell-lines, but immortalized cell-lines often lose their differentiation potential over time, it would be interesting to see if the LTS combo could extend the self-renewal of other tissue stem cells while preserving their differentiation potential, in future work.

Mechanistically, we found that *let-7* microRNAs could not fully explain Lin28a’s effects on muscle progenitors. Lin28a promoted the self-renewal of human myoblasts by optimizing OxPhos and mtROS, thereby inducing a series of mitohormetic stress-responsive pathways, including the HIF1A-mediated hypoxic response. Mitochondrial and hypoxic signatures were also observed in Lin28a+ MuSCs. This is interesting because multiple previous studies have shown that ROS activates HIF1A to drive the hypoxic stress response^63^. Moreover, it is well-known that hypoxia and HIF1A can transactivate glycolysis genes to promote proliferative capacity^63^. Previous findings have also shown that endogenous HIF1A is necessary for Notch signaling, muscle progenitor self-renewal, and myogenesis, but the upstream regulators and downstream effects on epigenetics had remained unclear^64^. Our findings now suggest that Lin28a-mtROS might be part of the upstream regulators of HIF1A during myogenesis and muscle regeneration, with dedifferentiation effects downstream. These results extend the emerging notion that, while excessive mtROS can cause a variety of degenerative phenotypes in many contexts, mild mtROS can be beneficial at the right amounts in the right cell-types (Sena and Chandel, 2012). Our study has provided a novel and tangible genetic combination that optimizes mitochondrial superoxides to rejuvenate stem cells’ biological age.

## Materials and Methods

### Transgenic mice

The targeting strategy for the generation of Lin28a-T2A-CreERT2 mice is shown in Figure S1A. With G418 selection, the targeted ES cell clones were selected. Next, the Frt-Neo-Frt expression cassette in selected ES cell clones was deleted and the resultant ES cells were injected into C57BL/6 albino embryos. The original animal was identified by its coat color, and its germline transmission was confirmed by reproduction with female C57BL/6 and subsequent genotyping. In the construction of the targeting vector to generate NFκB-LSL-Lin28a-T2A-luc (No. mt190) mice, the NF-κB response element and its downstream TAp promoter (the minimal TA promoter of the herpes simplex virus)^65^ were inserted upstream of a fragment of LSL. The targeting vector was integrated into the H11 site of C57BL/6 mice as previously reported to ensure the specificity of the NF-κB response element^66^. R26-tdTO ((*ROSA)26Sor^tm14^*^(*CAG*−*tdTomato*)^, stock no. 007914) was obtained from JAX laboratory. All animal procedures were approved by the Institute of Zoology and the Institute of Stem Cell and Regenerative Medicine, Chinese Academy of Sciences.

### Tamoxifen and cryoinjury treatment

Tamoxifen (TMX, Sigma-Aldrich) was dissolved at 20□mg□ml^−1^ in corn oil and 100mg/kg of TMX was administered by intraperitoneal injection into 6-week old mice, as schematized in the figures. For cryoinjury, all mice were anesthetized using isoflurane. The skin was cut open with a scalpel to expose the tibialis anterior (TA) muscle. A steel probe with a diameter of 4mm was cooled in liquid nitrogen and placed on the TA muscle for 10 seconds, twice. Subsequently, the skin incision was closed with surgical sutures immediately and betadine was applied to the wound to prevent infection after injury. The TA and soleus muscles were harvested 10 or 14 days after cryoinjury.

### Immunofluorescence

After skeletal muscles were harvested, they were fixed in 4% paraformaldehyde at 4□°C for 1□h. Tissues were then switched to 20% sucrose/ddH_2_O at 4□°C overnight, after this, they were freeze embedded with OCT and sectioned with 10μm-thickness. Before immunostaining, frozen slides were fixed in methanol for 10 minutes at room temperature and then washed with PBS. Next, slides immersed in citrate antigen retrieval solution (ab208572) were placed in a pressure cooker. Then, slides were treated with 0.3% Triton X-100/PBS at room temperature for 15 minutes, before incubation with mouse IgG-blocking solution (M.O.M. kit, Vector Lab) at room temperature for 1 hour, according to the manufacturer’s instructions. Slides were incubated with primary antibodies diluted in 0.3% Triton X-100, 5% goat serum/PBS at 4°C overnight. The next morning, slides were washed with PBS and incubated with secondary antibodies diluted in 0.3% Triton X-100, 5% goat serum/PBS at room temperature and protected from light for 1h. Slides were then washed with PBS and sealed with glycerine with DAPI. For immunofluorescence, the following antibodies were used: Pax7, Pax3, myosin I (BA-D5), myosin IIa (SC-71) (all from Developmental Studies Hybridoma Bank (DSHB), 1:10), IIx, IIb (Fast Myosin, Abcam, ab91506), laminin (Sigma-Aldrich, no. L9393, 1:500), desmin (abcam, ab32362, 1:200). Alexa Fluor secondary antibodies were used according to the manufacturer’s instructions. Images were taken on a Nikon confocal microscope or PerkinElmer Vectra Polaris. For muscle fiber type identification, type I and IIa myofibers were identified by antibodies against myosin I (BA-D5) and myosin IIa (SC-71) directly, type IIb myofibers were identified by light staining with Fast Myosin antibody (Abcam, ab91506), type IIx myofibers were identified by intense staining with Fast Myosin antibody (Abcam, ab91506) but absence of staining with myosin IIa antibody (SC-71).

### Fluorescence-activated cell sorting analysis

Conventional Pax7+ MuSCs were isolated by FACS sorting as the VCAM1 positive and CD45/CD31/Sca1-negative population, as described previously ^67^. Mononuclear cells were isolated as previously described ^68^. Following isolation, mononuclear cells from one mouse (~10^7^ cells) were resuspended in 500ul PBS/10% FBS/3mM EDTA, then incubated on ice for 40min with the following fluorophore-conjugated antibodies: APC-CD31 (clone MEC13.3), FITC-Sca1 (clone E13-161.7), VCAM1-biotin (clone 429) (all from Biolegend, 1:100), BV421-CD45 (Becton Dickinson, clone 30-F11, 1:250), and 20min with PE-Cy7 streptavidin (BioLegend, cat. no. 405206, 1:100). FMO controls were performed on these cells. Cells isolated from wild-type mice staining the same antibodies were used as the tdTomato FMO control. Unstained cells isolated from wild-type mice were used as unstained controls. Cells were analysed on BD FACSAria fusion flow cytometer and FACS data were analysed using FlowJo Software (TreeStar). Values of FACS analysis were averaged from more than three independent experiments.

### Cell culture and myogenesis, adipogenesis and osteogenesis *in vitro*

Young adult primary human skeletal muscle (HSKM) progenitors isolated from a 20-year old female subject’s quadriceps muscles (Gibco) were seeded onto plates coated with 0.1% gelatin solution (Merck-Millipore). Lin28a+ cells and conventional MuSCs cells were cultured in Matrigel-coated plates All cells were incubated at 37°C, 5% CO_2_ with growth medium (GM), comprising of DMEM/F-12 (Gibco) with 20% fetal bovine serum (FBS) (GE Healthcare), 1% L-glutamine (Gibco) and 1% penicillin-streptomycin (Gibco). At each passage, after reaching 80% confluence, cells were trypsinized and diluted 1:4. Differentiation was initiated by replacing growth media with differentiation medium (DM), comprising of DMEM/F-12, 2% KnockOut Serum Replacement (Gibco), 1% L-glutamine (Gibco) and 1% penicillin-streptomycin (Gibco), when the young adult HSKM progenitors were 80-100% confluent. These young adult HSKM progenitors were previously validated to be 100% MYOD1+ during the proliferative stage, with robust expression of myogenic markers after differentiation^40^. For fusion comparison experiments, conventional MuSCs stained with CellTrace Violet (Thermo fisher, C34557), Lin28a+ cells and 1:1 mixed cells were cultured in GM and then switched to DM to induce myotube formation, respectively. Freshly sorted Lin28a+ cells were grown in GM for 3 days and subsequently differentiated towards myotubes, adipocytes or osteoblasts. For myotube differentiation, the growth medium was replaced with DM for 2-3 days. Cells were then immunostained for MyoG (Santa Cruz Biotechnology, sc-12732, 1:100) and MYH1 (clone MF20, DSHB, 5μg/ml) to visualize differentiated myotubes. For adipocyte and osteoblast differentiation, cells were treated and detected by staining as previously described ^69^.

### Aged patient cells

Aged primary human skeletal muscle progenitors were derived from the rectus abdominus of aged patient donors with cachexia during tumour-resection surgery. Procedures were performed in accordance to ethical legislation and local Institutional Review Board guidelines. Muscle progenitors were isolated according to previously published protocols^28^. Cells were maintained in culture medium and differentiated with differentiation medium as described above.

### Immunostaining cultured cells

For immunostaining, cells grown on plates were fixed with 4% paraformaldehyde at room temperature for 15□min and subsequently permeabilized with 0.3% Triton X-100/ PBS. Cells were blocked in 10% goat serum diluted in 0.1% Triton X-100/ PBS at room temperature for 1□h. Primary antibodies were diluted in 1% goat serum/ 0.1% Triton X-100 and incubated at room temperature for 2 h. Cells were washed and secondary antibodies were diluted in 1% goat serum/0.1% Triton X-100 and incubated at room temperature for 1□h. Cells were stained with Hoechst dye (1:2,000 in PBS) at room temperature for 10 min. Primary antibodies include: Pax3 (DSHB, 1:20), Pax7, (DSHB, 1:20), MyoD (Santa Cruz Biotechnology, sc-377460, 1:100), MyoG (Santa Cruz Biotechnology, sc-12732, 1:100), MYH1 (clone MF20, DSHB, 5μg/ml). MYHC-IIb eFluor 660 (50-6503-32; Thermo Fisher; 1:100), α-actinin (sc-7453; Santa Cruz; 1:500), 8-oxo-guanine (ab206461; Abcam; 1:400). Fusion index was calculated as a ratio of the number of tdTO+ or MuSC nuclei within a multinucleated myotube to the total number of tdTO+ or MuSC nuclei. A minimum of 4 independent microscopic fields was used for each group over three independent differentiation experiments at 24 and 36 hours in DM.

### DNA/RNA isolation and NGS analysis

Cells were resuspended in 500ul of Trizol, genomic DNA and total RNA were isolated according to the manufacturer’s instructions (Invitrogen). RNA quality was verified by the Agilent 2100 Bioanalyzer with the RIN≥7, 28S/18S ≥1.5:1. RNA quantity was verified by QUBIT RNA ASSAY KIT. cDNA libraries were constructed using the NEBNext Ultra RNA Library Prep Kit for Illumina. All next generation sequencing (NGS) was performed using Illumina Novaseq-6000.

### Real-time Quantitative PCR analysis

Total RNA was extracted from sorted cells with Trizol (Invitrogen) following the manufacturer’s instructions. From this RNA, cDNA was reverse transcribed with PrimeScript RT reagent Kit (Takara, RR047B). The resulting cDNA was diluted 5x before performing qPCR with qPCR SYBR Green Mix. QPCR primer sequences were obtained from OriGene website. For miRNA quantitative PCR, 500 ng of total RNA extracted by TRIzol (Thermo Fisher) was amplified by miScript RT Kit (Qiagen). With the resulting cDNA libraries, real-time quantitative PCR was performed using the miScript SYBR Green PCR kit (Qiagen) on ABI Prism 7900HT (Applied Biosystems) according to manufacturers’ instructions. The following miScript Primer Assays (Qiagen) were used: Hs_let-7a_2 (MS00031220), Hs_let-7b_1 (MS00003122), Hs_let-7e_3 (MS00031227), Hs_let-7g_2 (MS00008337), Hs_RNU6-2_11 (MS00033740) and Hs_SNORD61_11 (MS00033705).

### *let-7* microRNA mimic

HSKM, TS and LTS cells were seeded in 6-well plates at a density of 4×10^4^ cells/well and maintained with culture medium. PEI (14.5 ng/μL), mature Mirvana hsa-let-7a-5p (#4464066; Assay ID: MC10050; Thermo Fisher) and/or hsa-let-7b-5p (#4464066; Assay ID: MC11050; Thermo Fisher) mimic (0.25 μM) or Cy5-conjugated scramble RNAi control, was mixed with serum-free DMEM to a total volume of 200 μL. This mixture was transfected onto each well.

### Western blot analysis

Protein was extracted with RIPA buffer supplemented with protease inhibitor cocktails I and II (Sigma) and phosphatase inhibitor cocktail set III (Calbiochem). Protein was quantified with Pierce BCA protein assay kit (Thermo Fisher) and analysed with a Sunrise Tecan plate reader. After SDS-PAGE and electro-transfer onto PVDF membranes, Western blotting was performed with the following primary antibodies and concentrations: Lin28a (1:1000, CST), Pax7 (0.28μg/ml, DSHB), Pax3 (0.31μg/ml, DSHB), MyoD (1:1000, Santa Cruz Biotechnology), MHC (0.23μg/ml, DSHB), MyoG (1:1000, Santa Cruz Biotechnology), IGF2BP2 (1:1000, Proteintech), Hmga2 (1:1000, CST), Cre (1:1000, Millipore), GAPDH (1:1000, CST), Vinculin (K106900P, 1:5000, Solarbio), Tubulin (ab210797; Abcam; 1:1000), P53 (sc-126; Santa Cruz; 1:100), IMP1/2/3 (sc-271785; Santa Cruz; 1:1000), citrate synthase (G-3) (sc-390693; Santa Cruz ; 1:1000).

### Virus production

The following plasmids were used to produce viruses for the various transgenic cell lines: lentiviral plasmid (Addgene #19119), dR8.2 packaging plasmid (Addgene #8455), VSV-G envelope plasmid (Addgene #8454). Viral supernatants were collected within the 48-hour to 96-hour window and filtered with a 0.45 μm filter (Sartorius).

### Population doubling curve

1.5×10^4^ cells were seeded in one well of a 6-well plate (Falcon) with growth medium comprising of DMEM/F-12 (Gibco) with 20% fetal bovine serum (FBS) (GE), 1% L-glutamine (Gibco) and 1% penicillin-streptomycin (Gibco). Upon reaching a confluency of 80-100%, cells were lifted with 0.25% trypsin (Gibco) and counted, 1.5×104 cells were then subcultured. This process was repeated until cells could no longer achieve 80% confluency, or until a period of 100 days. Recorded cell counts were calculated as cumulative population doubling levels and plotted over the number of days in culture.

### Senescence-associated β-galactosidase assays

Senescence-associated β-galactosidase (SA-β-gal) activity was determined with the Senescence Cells Histochemical Staining Kit (Sigma-Aldrich). Six representative images were captured on a TS100 inverted light microscope (Nikon) and camera module. For each cell line, cells stained blue by cleaved X-gal, indicating the presence of SA-β-gal, were counted as a percentage of the total number of cells in all images.

### Cytogenetics

Cells were seeded in gelatin-coated six-well plates and cultured up to a maximum of 50% confluency to avoid myoblast fusion. Cells were first treated with colcemid and BrdU overnight before harvesting with EDTA. A fixative solution (1:3=glacial acetic acid:methanol) was used to fix pelleted cells prior to slide preparation, Giemsa banding and mounting. Twenty metaphase spreads were prepared for each cell line for detailed analyses and karyotyping.

### Extracellular flux and oxygen consumption measurements

Cells were seeded onto Seahorse XFe96 cell culture plate (Agilent) at a density of 6000 - 8000 cells/well with culture medium for 24 hours. Subsequently, media was changed to Seahorse minimal media and incubated in a non-CO2, 37°C incubator for one hour as per manufacturer’s instructions. Utility plate was loaded with the following nutrient/ drugs, Port A – 10 mM glucose (for glycolysis) or 10 mM pyruvate (for OxPhos); Port B– 1 μM oligomycin; Port C– 0.5 μM FCCP; Port D– 0.5 μM antimycin A. Assay was run on a Seahorse XF extracellular flux analyser (Agilent) according to manufacturer’s protocols.

### KC7F2, ROS modulators and 2-deoxyglucose (2-DG)

HSKM, TS and LTS cells were seeded in gelatin-coated six-well plates at a density of 4×104 cells/well and maintained with culture medium. One day after seeding, culture medium was replaced with culture medium containing either paraquat (Sigma-Aldrich; 2 μM), hydrogen peroxide (ICM Pharma; 100 μM), DTT (Thermo Fisher; 250 μM), 2-deoxyglucose (Sigma-Aldrich; 2.5 mM), KC7F2 (Sigma-Aldrich; 10μM) or DMSO/ H2O controls. For ROS modulators, cells were allowed to grow for 72 hours post-treatment, while for 2-DG, cells were allowed to grow for eight days post-treatment.

### Flow cytometry assay for oxidation state with mito-Grx1-roGFP2

Retroviruses were created with the pLPCX mito Grx1-roGFP2 plasmid according to the virus production subsection. HSKM, TS and LTS cells were transduced with the roGFP2 retrovirus for 48 hours. Cells were allowed to expand to 15-cm tissue culture plates and then harvested for fluorescence-activated cell sorting for roGFP2+ cells using the FITC channel with a BD FACS Aria II cell sorter. HSKM, TS and LTS cells were seeded in gelatin-coated six-well plates at a density of 4×104 cells/well and maintained with culture medium. One day after seeding, culture medium was replaced with culture medium containing either paraquat (Sigma-Aldrich; 2 μM), hydrogen peroxide (ICM Pharma; 100 μM), or DTT (Thermo Fisher; 250 μM). Cells were allowed to grow for 48 hours and then harvested for flow cytometry using a BD LSR Fortessa x-20 analyzer. HSKM, TS and LTS cells were used as non-fluorescent gating controls for each of their respective roGFP2+ counterparts. For each cell line, FSC, SSC, 405 nm/AmCyan and 488 nm/FITC channels were captured. FCS files were exported from FACS Diva and imported into FlowJo, where raw-value CSV files were exported. Raw values of 405 nm/488 nm ratio was calculated and then averaged for each cell line and condition. The % oxidised value was calculated according to previously published methods ^70^, where the difference between the 405/488 value of the vehicle control and DTT-treated condition was divided by the difference between the 405/488 value of the hydrogen peroxide and DTT-treated conditions.

### Statistics and reproducibility

All statistical analyses were performed using GraphPad Prism 6 (GraphPadSoftware). Data are presented as mean ± s.e.m. Differences between groups were tested for statistical significance by using the two-sample *t*-test. *P* < 0.05 was considered significant. The number of biological (non-technical) replicates for each experiment is indicated in the figure legends.

## Author contributions

P.W., X.L. and J.H.E.T. designed and performed the experiments. P.W., X.L., J.H.E.T. and N.S.-C. wrote the manuscript. L.G. helped with various experiments and mouse husbandry. J.H.E.T., M.W.J.C. and Y.J.B.C. helped with HSKM and LTS cell lines experiments. L.L. helped with cryosection and immunofluorescence experiments. S.M. helped with Western Blot experiments. W.C. helped with quantitative RT-PCR experiments. W.M. and A.N. helped with genomics and bioinformatics analysis. Z.Y., Y.C., H.M., L.G., K.L., Y.W., J.S., R.R.W., C.L., N.A., Y.L., Z.J., T.L. provided technical assistance. L.Z., Z,J. and T.L. helped with mouse sampling. B.T.T., L.J., K.Y., and N.S.-C. designed and supervised the entire project.

## Acknowledgements

This work was supported by the Strategic Priority Research Program of the CAS (XDA16010109), the National Key R&D Program of China (2019YFA0801701, 2019YFA0801702, 2018YFE0201100, 2018YFC1004102), the National Natural Science Foundation of China (1191957202, 81900782), the Key Research Program of the CAS (KJZD-SW-L04), and the State Key Laboratory of Stem Cell and Reproductive Biology. N.S.-C. is also a Howard Hughes Medical Institute (HHMI) International Scholar.

---

**Figure S1.**
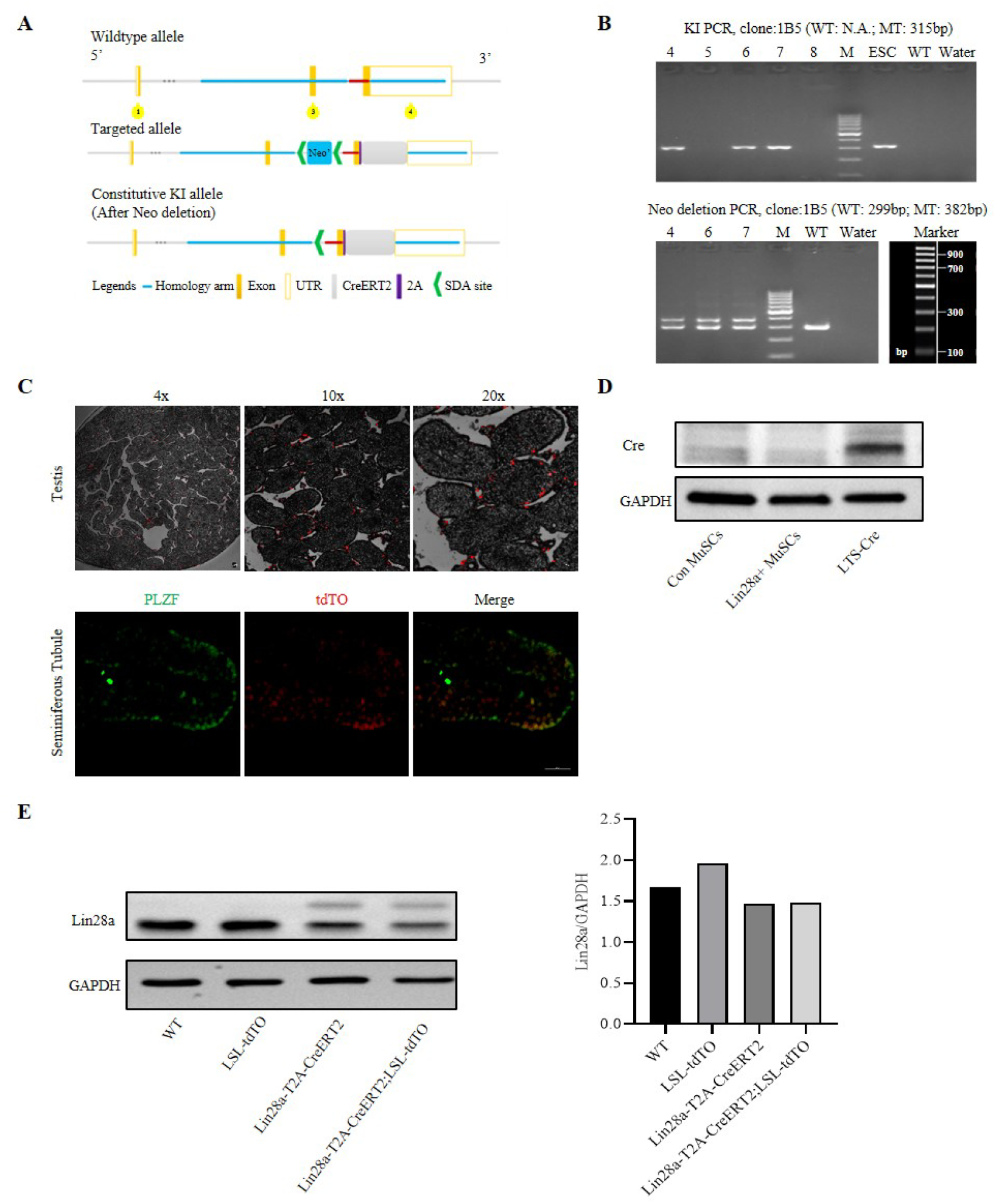
Generation of Lin28a-T2A-CreERT2 mice and evaluation of lineage tracing. **(A)** The targeting strategy for the generation of Lin28a-T2A-CreERT2 mice. The fragment of CreERT2 was inserted between the last exon and the 3’UTR of Lin28a. **(B)** Genotyping results of Lin28a-T2A-CreERT2 mice. **(C)** Lin28a-T2A-CreER;LSL-tdTO mice’ testis shows a subset of PLZF+ spermatogonial stem cells (SSCs) are tdTO+ after 14 days of lineage tracing. Scale bar 50 μm. **(D)** Western blot analysis for Cre protein in conventional (Con) MuSCs and tdTO+ MuSCs shortly after culture in vitro. Lin28-Cre cells are positive control cells which overexpressed Cre protein. Lin28a expression is extinguished shortly after Con MuSCs and tdTO+ MuSCs were cultured in vitro (Fig S3A). **(E)** Western blot analysis for Lin28a protein in the testis of wildtype (WT), LSL-tdTO, Lin28a-T2A-CreERT2 and Lin28a-T2A-CreER;LSL-tdTO mice respectively. Lin28a-T2A-CreERT2 mice manifested two bands: the lower band of endogenous Lin28a and the higher band of Lin28a-T2A. Both bands’ intensities were quantified for the Lin28a/GAPDH ratio. GAPDH protein was used as the loading control.

**Figure S2.**
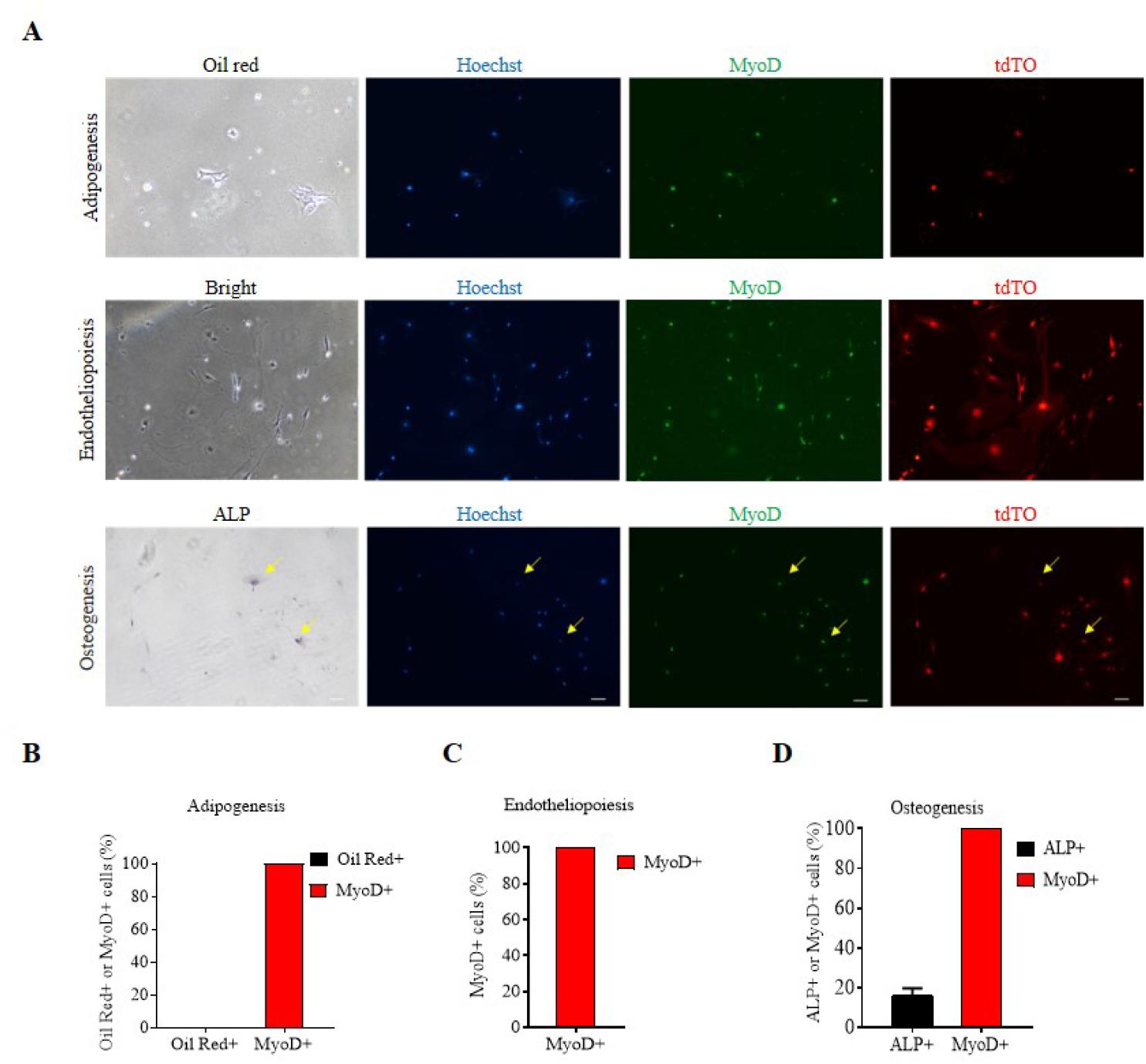
In vitro differentiation of freshly sorted Lin28a+ MuSCs. **(A)** Phase contrast and fluorescence microscopy to assess Lin28a+ cells after in vitro differentiation in adipogenic, endothelial or osteogenic conditions. Alkaline phosphatase (ALP) staining revealed that, after differentiation in osteogenic conditions for 10 days, tdTO+ cells still expressed MyoD. The yellow arrows indicate some tdTO+ cells that are lightly stained by ALP and co-stain for MyoD. Scale bar: 100 um. **(B)** Proportion of Oil Red-positive cells and MyoD-positive cells per field after culture in adipogenic conditions. Data are mean ± SEM. N =3 independent experiments. For each experiment, total of 5 fields were counted and averaged. **(C)** Proportion of MyoD-positive cells per field after culture in endothelial conditions. Data are mean ± SEM. N =3 independent experiments. For each experiment, total of 5 fields were counted and averaged. **(D)** Proportion of ALP-positive cells and MyoD-positive cells per field after culture in osteogenic conditions. Data are mean ± SEM. N =3 independent experiments. For each experiment, total of 5 fields were counted and averaged. ** P < 0.01.

**Figure S3.**
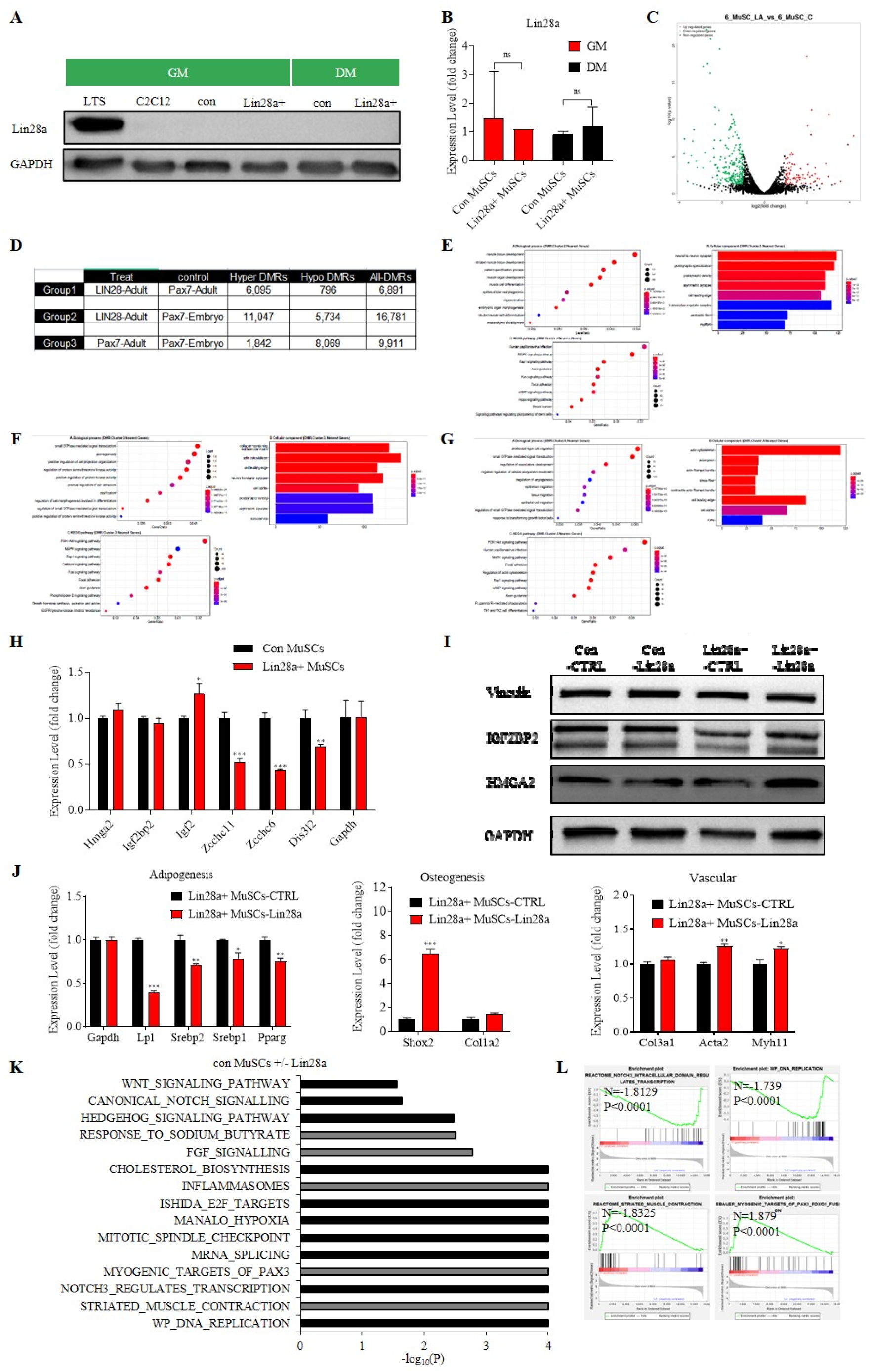
Gene expression in conventional and Lin28a+ MuSCs. **(A)** Western blot analysis for Lin28a protein in mouse C2C12 myoblasts, conventional (con) MuSCs and Lin28a+ cells in myogenic growth medium (GM) and differentiation medium (DM). LTS is a positive control cell-line which overexpressed Lin28a. Lin28a expression was extinguished once cells were plated in culture, and it was not detected in both GM and DM. GAPDH protein was used as the loading control. **(B)** Quantitative RT-PCR revealed expression levels of Lin28a in conventional (con) MuSCs and Lin28a+ cells in myogenic growth medium (GM) and differentiation medium (DM). Data are mean ± SEM. N = 3 independent experiments. **(C)** Volcano plot analysis for genes that are differentially expressed in conventional MuSCs overexpressing Lin28a, relative to conventional MuSCs. **(D)** Whole-genome bisulfite sequencing (WGBS) analysis for Lin28a+ MuSCs to embryonic and adult Pax7+ MuSCs (N=3 mice each). **(E)** GO and KEGG analysis of Cluster 2 in Figure 3A. **(F)** GO and KEGG analysis of Cluster 3 in Figure 3A. **(G)** GO and KEGG analysis of Cluster 5 in Figure 3A. **(H)** Quantitative RT-PCR analysis of *let-7* targets (*Hmga2, Igfbp2*), *Igf2,* and *let-7* pathway-associated genes (*Zcchc6, Zcchc11, Dis3l2*) expression in freshly sorted con MuSCs and Lin28a+ MuSCs in GM. **(I)** Western blot analysis for Lin28a and *let-7* targets IGF2BP2 and HMGA2 in con MuSCs or Lin28a-tdTO+ MuSCs which overexpressed Lin28a, relative to con MuSCs or Lin28a-tdTO+ MuSCs with the empty vector (CTRL). **(J)** Quantitative RT-PCR for adipogenesis, osteogenesis and vascular genes expression in Lin28a-tdTO+ MuSCs overexpressing Lin28a relative to the empty vector (CTRL). Data are mean ± SEM. N = 3 independent experiments. **(K)** Signatures enriched in con MuSCs overexpressing Lin28a (black) or empty vector (grey), as identified by Gene Set Enrichment Analysis. **(L)** Representative GSEA profiles with normalized enrichment scores (NES) and nominal P values are shown. Ns, not significant. * P < 0.05, ** P < 0.01, *** P < 0.001.

**Figure S4.**
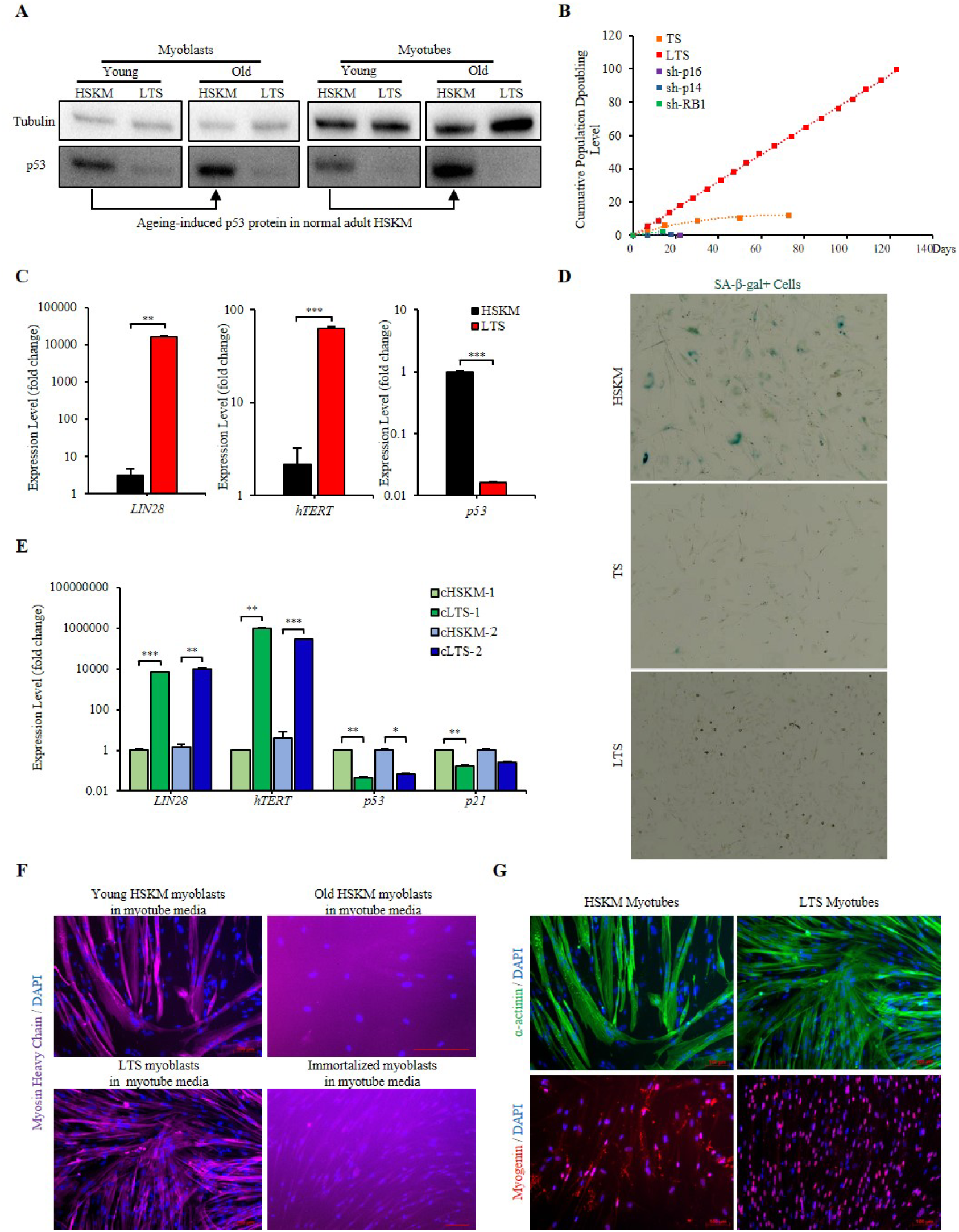
Senescence, self-renewal and differentiation capacities of LTS and HSKM human progenitors. **(A)** Western blot analysis of P53 protein, relative to tubulin protein, in young and aged adult HSKM and LTS myoblasts and myotubes. **(B)** Population doubling curves for young HSKM and other transgenic myoblasts. Young adult HSKM myoblasts transduced with hTERT and sh-p53 (TS, orange), or Lin28a, hTERT and sh-p53 (LTS, red) showed different proliferation rates. Lentiviral shRNA knockdowns of *P16^INK4A^* (sh-p16, purple), *P14^ARF^* (sh-p14, blue), and *RB1* (sh-RB1, green) were also attempted, but were not followed up upon as they appeared to be prematurely senescent. **(C)** Quantitative RT-PCR for mRNAs of Lin28a, *hTERT*, and *TP53* in LTS myoblasts, relative to young adult HSKM myoblasts. **(D)** Brightfield micrographs of senescence-associated β-galactosidase positive (SA-β-gal+) cells in 100-day-old adult HSKM myoblasts and 100-day-old LTS myoblasts. **(E)** Immunofluorescence staining for the myotube protein marker myosin heavy chain (MHC) in young HSKM and LTS myoblasts, relative to old HSKM myoblasts and immortalized (hTERT-CyclinD1-CDK4^R24C^) myoblasts, that were cultured in myogenic differentiation media. Cells were counterstained with DAPI to visualize the myonuclei. Scale bars 100 μm. **(F)** Immunofluorescence staining for the myotube protein markers α-actinin (green) and nuclear myogenin (red) in young HSKM myotubes, relative to LTS myotubes. Cells were counterstained with DAPI (blue) to visualize the myonuclei. Scale bars 100 μm. * P < 0.05, ** P < 0.01, *** P < 0.001.

**Figure S5.**
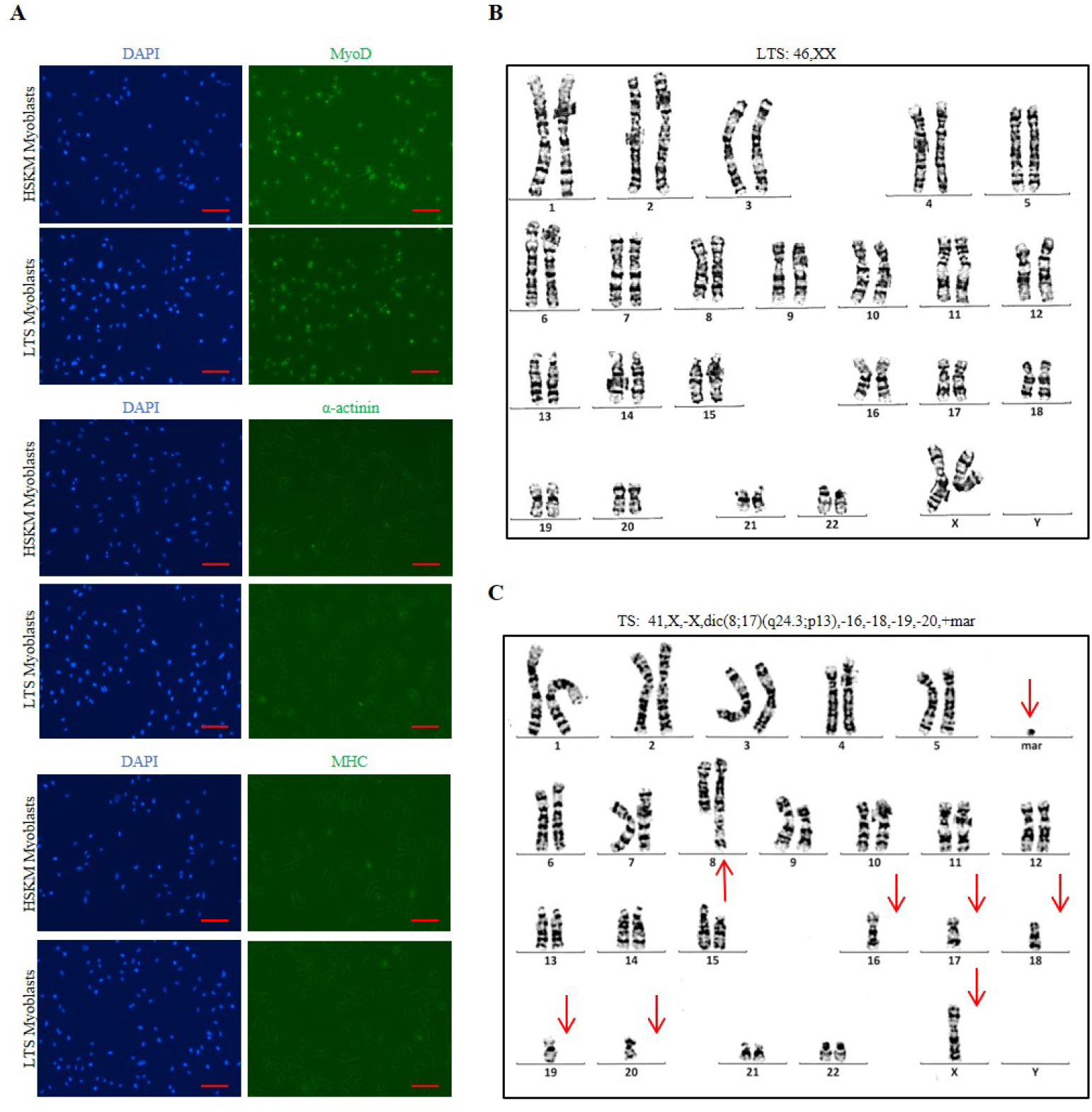
Protein expression and karyotype analysis of LTS human progenitors. **(A)** Immunofluorescence staining for the myogenic protein markers MyoD, α-actinin, and myosin heavy chain MHC (green) in young LTS myoblasts, relative to HSKM myoblasts. Cells were counterstained with DAPI (blue) to visualize the myonuclei. Scale bars 100 μm. (**B**-**C**) Representative karyotype analysis of old LTS (**B**) and TS (**C**) myoblasts, which are [46,XX] and [41,X,-X,dic(8;17)(q24.3;p13),-16,-18,-19,-20,+mar], respectively. Chromosomal aberrations are indicated with red arrows.

**Figure S6.**
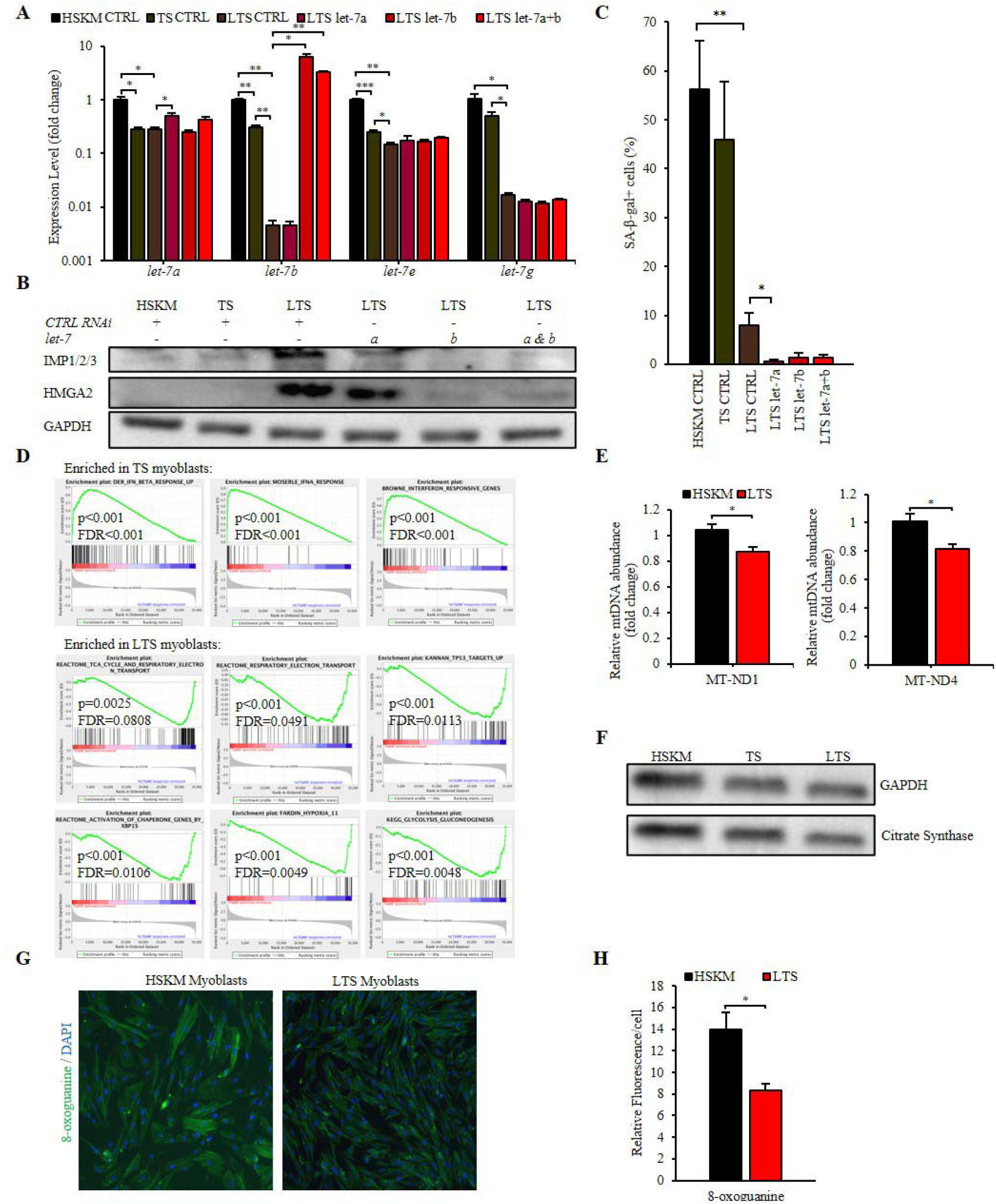
The mechanism underlying LTS progenitors’ self-renewal. **(A)** Quantitative RT-PCR for *let-7* microRNAs in TS and LTS myoblasts, relative to young adult HSKM myoblasts, after control (CTRL), *let-*7a, *let-7b,* or combined *let-*7a and *let-7b* (let-7a+b) mimic transfection. **(B)** Western blot analysis for the *let-7* targets IGF2BP1/2/3 (IMP1/2/3) and HMGA2 protein, relative to GAPDH protein, in young adult HSKM, TS and LTS myoblasts, after control (CTRL), *let-*7a (a), *let-7b* (b), or combined *let-*7a and *let-7b* (a & b) mimic transfection. **(C)** Quantification of SA-β-gal+ cells in TS and LTS myoblasts, relative to young adult HSKM myoblasts, after control RNAi (Ctrl RNAi), *let-*7a, *let-7b,* or combined *let-*7a and *let-7b* (let-7a+b) mimic transfection. **(D)** Representative GSEA enrichment profiles for RNAseq analysis of TS and LTS myoblasts. **(E)** Quantitative RT-PCR for mitochondrial NADH-ubiquinone oxidoreductase chain 1/4 (MT-ND1/4), relative to the nuclear reference gene B2M, to assess mtDNA copy number in adult HSKM myoblasts, relative to LTS myoblasts. **(F)** Western blot analysis for mitochondrial citrate synthase protein, relative to GAPDH protein, in TS and LTS myoblasts, relative to adult HSKM myoblasts. **(G)** Immunofluorescence staining for the DNA oxidative damage biomarker 8-oxo-guanine (green) in adult HSKM and LTS myoblasts. Cells were counterstained with DAPI (blue) to visualize the myonuclei. **(H)** Quantification of the immunofluorescence signal from 8-oxo-guanine in adult HSKM and LTS myoblasts. * P < 0.05, ** P < 0.01.

**Figure S7.**
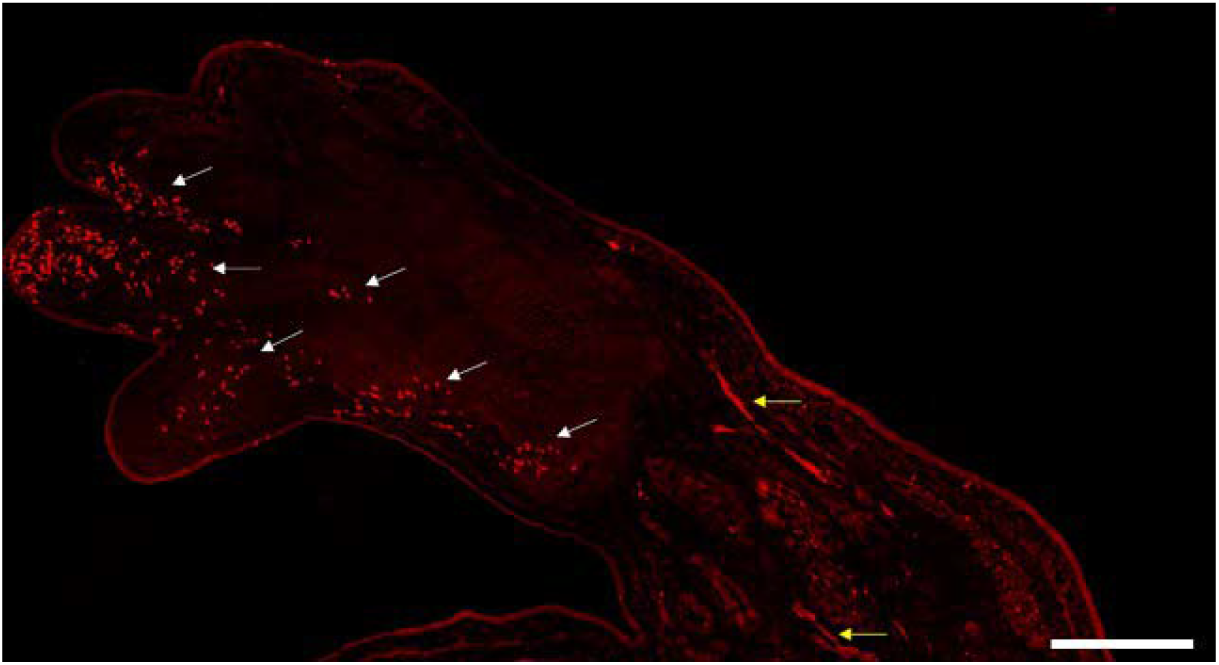
Lin28a+ cells in the limb bud of E12.5 embryos. Lin28a-T2A-CreER;LSL-tdTO embryos were treated with TMX at E11.5 and harvested at E12.5. White arrows indicate Lin28a+ limb progenitor cells, while yellow arrows indicate Lin28a+ myofibers in the embryonic limb. Scale bar: 500um. E12.5 = embryonic day 12.5 post coitum.

